# Understanding indirect assortative mating and its intergenerational consequences for educational attainment

**DOI:** 10.1101/2024.06.21.600029

**Authors:** Hans Fredrik Sunde, Espen Moen Eilertsen, Fartein Ask Torvik

## Abstract

We develop a framework for understanding indirect assortative mating and provide updated definitions of key terms. We then develop family models that use partners of twins and siblings to freely estimate the degree of genetic and social homogamy, and account for it when investigating sources of parent-offspring similarity. We applied the models to educational attainment using 1,545,444 individuals in 212,070 extended families in the Norwegian population and Norwegian Twin Registry. Partner similarity in education was better explained by indirect assortment than direct assortment on observed educational attainment, with social homogamy being particularly important. The implied genotypic partner correlation (*r*=.34) was comparable to earlier studies, and higher than expected under direct assortment. About 38% of the parent-offspring correlation (*r*=.34) was attributable to various forms of environmental transmission. Alternative models that assumed direct assortment estimated environmental transmission to be lower, but these did not fit the data well.

## Introduction

Mating partners tend to have similar educational attainment^1^. Researchers broadly agree that this results from assortative mating not only on education itself, but also on associated traits, as partner resemblance in factors relevant for educational attainment is estimated to be higher than expected given resemblance in observed educational attainment^2–9^. However, how the partner correlations arise remains a matter of debate. Understanding factors that contribute to partnership formation is crucial for a nuanced understanding of the mechanisms that contribute to social inequality within and across generations. It is also crucial for improving the validity of statistical models, in particular genetic models of intergenerational resemblance, which are biased in the presence of unmodelled or inaccurately modelled assortative mating^10–16^. Despite this, progress has been hampered by underdeveloped theories of indirect assortative mating and inconsistent use of relevant terms, which in turn have led to idiosyncratic modelling decisions and studies with disparate conclusions. In this paper, we provide a framework for understanding partner similarity and various forms of assortative mating. In so doing, we clarify the inconsistent terminology found in the literature and introduce refined definitions of terms that describe partner similarity. We then show how partners of twins can inform us on the nature of assortative mating and apply this on 212,070 extended families in Norwegian registry data to understand partner similarity in educational attainment and its intergenerational consequences.

### What is direct and indirect assortative mating?

Assortative mating arises when individuals with similar trait values mate more often than expected by chance. More specifically, it is when partner formation is partly conditional on individuals’ having similar trait values, resulting in covariance between partners’ traits. Because the trait values themselves remain unchanged, the variance is also unchanged. A defining feature of assortative mating as opposed to other causes of partner similarity is therefore that it induces covariance without affecting variance. Finally, assortative mating is typically positive (i.e., like mating with like). It can in principle be negative (often called disassortative mating), but negative correlations between partners are rare^1^. In this paper, we will assume assortment means positive assortment unless otherwise stated.

When mating is conditional on specific traits, the causes of these traits (and other associated factors) will become correlated across partners. For example, if the traits are associated with genetic differences, partners will tend to be genetically similar. Numerous studies have demonstrated genetic similarity between partners for traits like educational attainment^6,7,17-20^. Genetic similarity between partners will have numerous consequences in subsequent generations, including long-range linkage disequilibrium between trait-associated loci, increased genetic variance in the population, increased genetic similarity between relatives, and, for traits influenced by few loci, increased homozygosity at trait-associated loci^18,21-26^. This can bias estimates of genetic and environmental effects unless accounted for properly^12,14^. The magnitude of these consequences (and hence the potential biases) is a function of the genotypic correlation between partners. Because this correlation is often unknown, it is typically inferred based on the strong assumption that assortment takes place directly on the observed phenotype – henceforth called ‘the focal phenotype’ (**Figure 1a**). This is called direct assortment, or primary phenotypic assortment^27^. Under direct assortment, the expected genotypic correlation between partners can easily be calculated, and the consequences and necessary adjustments are well understood (see **Supplementary Note 1**). However, direct assortment on one phenotype will also lead to partner correlations on associated phenotypes^28^. For example, partners assorting on height will have correlated arm lengths as well^24^. These secondary partner correlations are said to result from indirect assortment, or secondary assortative mating^7^. Crucially, if partner correlations in the focal phenotype results from indirect assortment, the consequences will not follow the same expectations as for direct assortment and models can become biased if they incorrectly assume direct assortment^21^.

**Figure 1.**
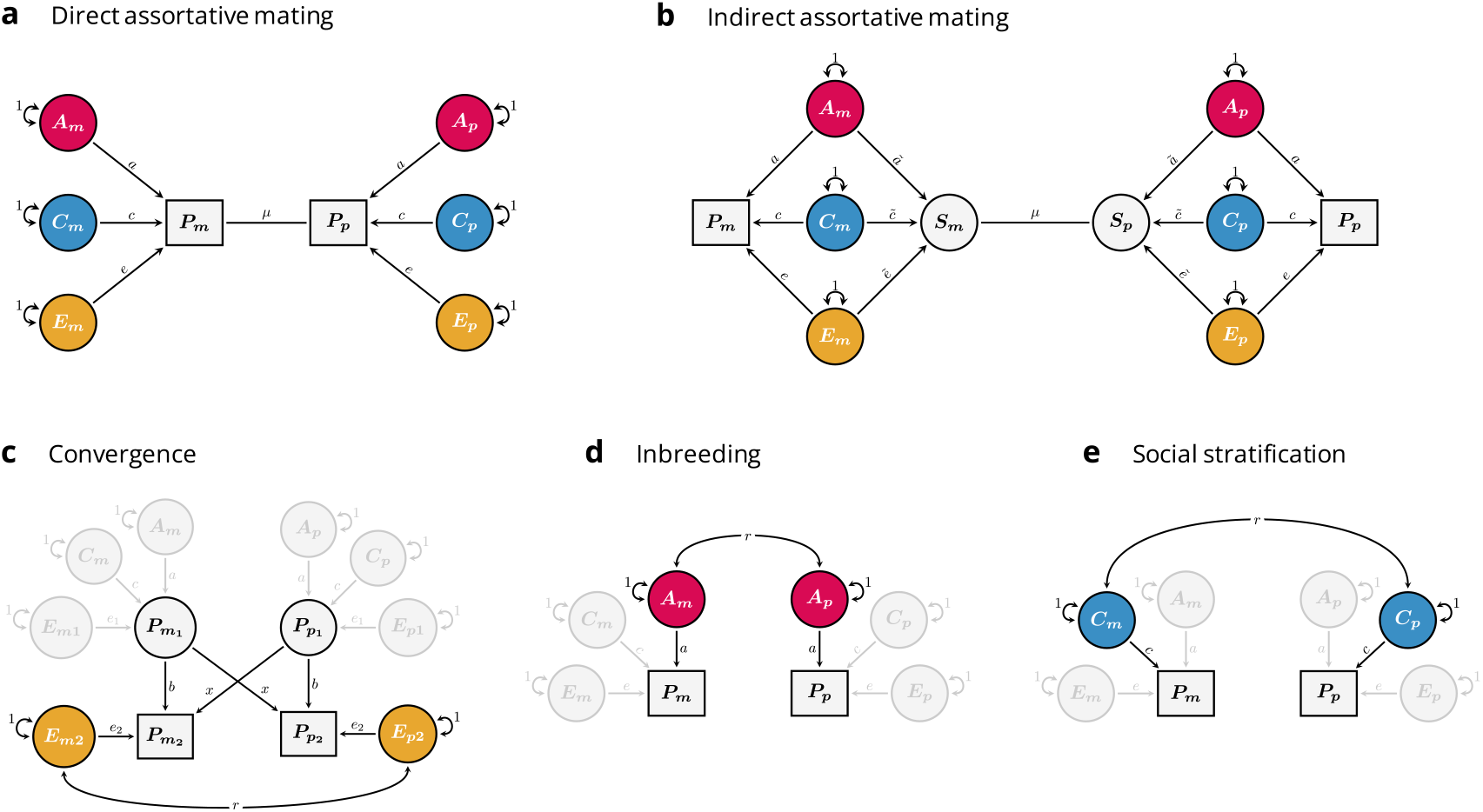
Causes of partner similarity illustrated with path models. Each panel includes two observed, focal phenotypes (rectangles denoted *P*), which belong to the two members of a partnership (*m* and *P*). *A, C*, and *E* refer to additive genetic (red), sibling-shared environmental (blue) and non-shared environmental influences (yellow) on the phenotype, respectively, whereas the different path coefficients (e.g., *a, c*, and *e*) refer to their effects. For simplicity’s sake, we assume no correlations between genetic and environmental influences, but this can be added to the models when relevant (see also Figure 5). The copath (the arrowless path denoted μ) denotes assortative mating on the two connected variables, which results in covariance without affecting variances. (**a**) Direct assortative mating: Here, partners are assorting directly on the focal phenotype, CoV(*P_m_, P_P_*) = μ(*a*^2^ + *c*^2^ + *e*^2^)^2^. (**b**) Indirect assortative mating: Here, the effects of *A, C*, and *E* on factors that are assorted upon (a sorting factor, denoted *S*) are allowed to differ from their respective effects on the focal phenotype, 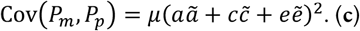Convergence: Here, partners become more similar over time, either due to mutual influence (denoted *x*) or exposure to similar environments (denoted *e*_2_), 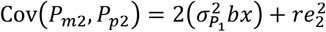, where 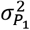 refers to the phenotypic variance prior to partner formation. (**d**) Inbreeding: Here, the two partner’s genetic effects are correlated, CoV(*P_m_, P_P_*) = *ra*^2^. (**e**) Social stratification: Here, the two partner’s sibling-shared environmental effects are correlated, CoV(*P_m_, P_P_*) = *rc*^2^.

### Evidence against direct assortment on educational attainment

Numerous lines of evidence suggest that direct assortment is an inadequate explanation for partner similarity in educational attainment, and that partners must instead be more similar on variables associated with education. One line of evidence comes from studies looking at trait-specific genetic similarity between partners. Robinson, et al. ^6^ found that education-associated genetic similarity between partners in the UK Biobank implied a phenotypic correlation of .65, much higher than the observed phenotypic correlation at .41. Similarly, in the latest genome-wide association study of educational attainment, Okbay, et al. ^8^ found that the polygenic index correlation between partners remained substantial even after adjusting for observed educational attainment and other indicators of cognitive performance. Finally, Torvik, et al. ^7^ reported a polygenic index correlation between partners that was too high given direct assortment. They expanded upon this by using genetic and phenotypic similarity between partners, siblings, and siblings-in-law and estimated a phenotypic partner correlation of .68 on an unobserved factor associated with educational attainment, with a genotypic correlation between partners estimated at .37.

Other studies have reached similar conclusions using phenotypic correlations between distant relatives, which increase under assortative mating on heritable traits. For example, Kemper, et al. ^29^ inferred the phenotypic correlation between spouses’ educational attainment by investigating phenotypic correlations between distant relatives in the UK Biobank. They concluded it must be higher than empirically reported elsewhere (.60 versus .42), suggesting indirect assortment. Similarly, Clark ^4^ investigated correlations in social class (a phenotype closely associated with educational attainment) among distant relatives in another large, English data set and found that genetic similarity alone could account the familial resemblance if it were the case that partners correlated .79 on an unobserved factor associated with social class, and this factor was highly heritable. On the other hand, Collado, et al. ^3^ investigated educational attainment among distant family members in Sweden by chain-linking affine kinships (i.e., in-laws) and, while also finding that partners must be highly correlated (.75) on an unobserved factor associated with educational attainment, they ascribed this largely to assortment on non-genetic factors. Twin studies that have included partners also conclude that partners are more environmentally similar than expected given direct assortment. A recent study of Finnish and Dutch twins and their spouses estimated that a third of the partner correlation could be attributed to so-called social homogamy (see below for a critical discussion about this term) ^5^. An earlier study by Reynolds, et al. ^2^ reported similar results for both educational attainment and cognitive performance in Swedish twins.

## Hindrances to understanding of indirect assortative mating

Despite converging evidence that partner similarity in educational attainment results from some form of indirect assortative mating, prior studies reach disparate conclusions about the nature of the traits partners are assorting on, ranging from the highly genetic to the highly cultural or environmental. One reason for the disparate findings, we believe, is an inadequate theoretical framework for indirect assortative mating. Previously described alternatives to direct assortative mating – such as genetic and social homogamy – are either underdefined or overly simplistic^7,19,27,30,31^. For example, social homogamy is often implied to mean assortment on environmental factors only^27,32^. As a result, many studies that have attempted to account for assortative mating have either assumed direct assortment^33,34^, or at best compared the fit of models assuming direct assortment with models assuming pure social homogamy ^35,36^. Attempts at more sophisticated models are often idiosyncratic and unsuitable. For example, many of the interpretations described above can be ascribed to the way the models have been specified rather than the way the world works. For example, Clark ^4^ did not model environmental similarity between family members, and thereby precluded a cultural explanation for partner similarity. While the model fits the data well, it is known that environmental and genetic transmission can give similar correlations in extended families and thereby equal fit^37^. Conversely, the models in Collado, et al. ^3^ and Reynolds, et al. ^2^ do not allow the implied genotypic correlation between partners to exceed expectations under direct assortment. Any deviations from direct assortment must therefore be attributed to assortment on cultural factors in these models.

One hindrance is inconsistent terminology. In particular, the term ‘social homogamy’ has been used inconsistently. Sometimes, it has been used to mean indirect assortative mating while remaining agnostic about the nature of the sorting process^30^. Most often, social homogamy is used to describe partner similarity arising for environmental reasons, but it is not consistent whether it refers to assortment on environmental factors or if it refers to shared environmental causes unrelated to assortment^2,5,6,17,27,31,38,39^ (see “other causes of partner similarity” below). Genetic homogamy is somewhat more clearly defined – assortment on genetic factors – although there remains some ambiguity as to whether social homogamy and genetic homogamy are meant to be mutually exclusive, and whether genetic homogamy is to mean assortment on the genotype^23,24^ (as if environmental influences on the phenotype were incidental) versus assortment on a sorting factor that is more heritable than the focal phenotype^1,12^. This inconsistency breeds confusion and hampers methodological and theoretical progress. For the sake of clarity and to foster a common understanding moving forward, we need more rigorous terms grounded in a coherent theoretical understanding of the sorting process.

### Defining genetic and social homogamy

The consequences of assortative mating, with respect to the focal phenotype, will depend on the magnitude of the induced correlation in causes, which in turn will depend on why the focal phenotype is correlated with factors that are assorted upon (see **Supplementary Note 1**). For example, the genetic consequences only depend on how much the genetic influences on the focal phenotype are assorted upon, regardless of whether assortment is direct or not. We therefore suggest defining genetic and social homogamy to reflect the degree to which genetic and social influences on the focal phenotype are associated with the factors that are assorted upon, henceforth collectively referred to as ‘the sorting factor’ (see **Table 1**). The sorting factor for a given phenotype refers to the associated trait or set of traits undergoing assortative mating and is indicated with the latent variable *S* in **Figure 1b**. Note that the sorting factor only includes the components of the associated traits that are associated with the focal phenotype (see **Supplementary Note 1**).

**Table 1.**
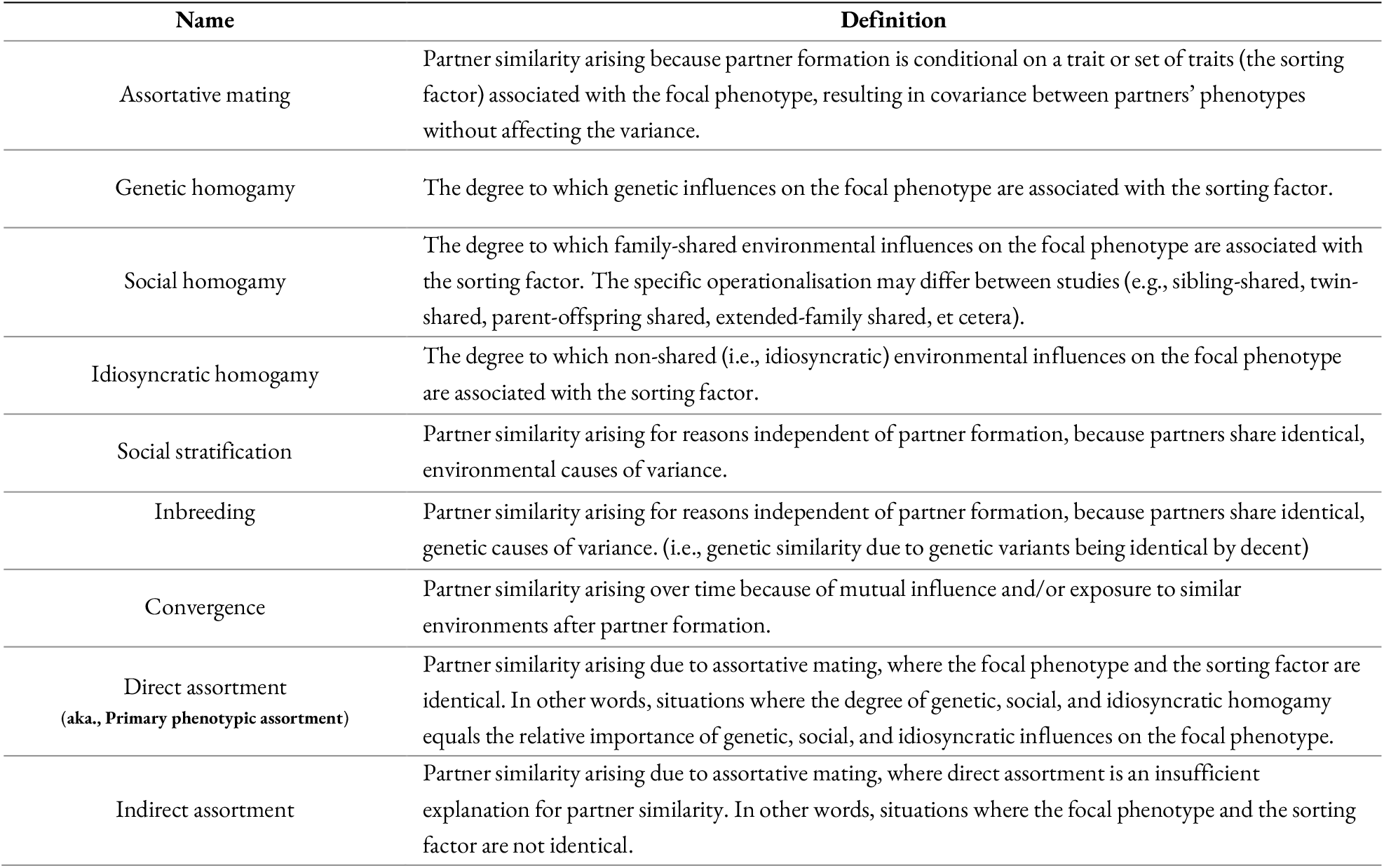
Causes of partner similarity.

| Name | Definition |
| --- | --- |
| Assortative mating | Partner similarity arising because partner formation is conditional on a trait or set of traits (the sorting factor) associated with the focal phenotype, resulting in covariance between partners' phenotypes without affecting the variance. |
| Genetic homogamy | The degree to which genetic influences on the focal phenotype are associated with the sorting factor. |
| Social homogamy | The degree to which family-shared environmental influences on the focal phenotype are associated with the sorting factor. The specific operationalisation may differ between studies (e.g., sibling-shared, twin-shared, parent-offspring shared, extended-family shared, et cetera). |
| Idiosyncratic homogamy | The degree to which non-shared (i.e., idiosyncratic) environmental influences on the focal phenotype are associated with the sorting factor. |
| Social stratification | Partner similarity arising for reasons independent of partner formation, because partners share identical, environmental causes of variance. |
| Inbreeding | Partner similarity arising for reasons independent of partner formation, because partners share identical, genetic causes of variance. (i.e., genetic similarity due to genetic variants being identical by decent) |
| Convergence | Partner similarity arising over time because of mutual influence and/or exposure to similar environments after partner formation. |
| Direct assortment<br>(aka., Primary phenotypic assortment) | Partner similarity arising due to assortative mating, where the focal phenotype and the sorting factor are identical. In other words, situations where the degree of genetic, social, and idiosyncratic homogamy equals the relative importance of genetic, social, and idiosyncratic influences on the focal phenotype. |
| Indirect assortment | Partner similarity arising due to assortative mating, where direct assortment is an insufficient explanation for partner similarity. In other words, situations where the focal phenotype and the sorting factor are not identical. |

Using this framework, we suggest defining genetic homogamy as the degree to which the genetic influences on the focal phenotype are also associated with the sorting factor (ã in **Figure 1b**). For traits influenced by both additive and non-additive (e.g., dominant) genetic effects, it can be useful to distinguish narrow-sense sense (additive only) and broad-sense (additive and non-additive) genetic homogamy. We urge readers to be explicit if they model varied forms of genetic homogamy. In this paper, we will only consider narrow-sense genetic homogamy.

As for environmental correlations between the focal phenotype and the sorting factor, it will be advantageous to separate environmental factors shared by family members and environmental factors unique to the individual (analogous to shared and non-shared environmental factors in the classical twin model). We suggest defining social homogamy as the degree to which family-shared environmental influences on the focal phenotype are associated with the sorting factor (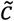 in **Figure 1b**). The specific operationalisation of family-shared environments may differ between studies (e.g., sibling-shared, twin-shared, parent-offspring shared, extended-family shared, et cetera), but we do not deem it worthwhile to define separate terms for all these contexts as their definition would simply follow how they are defined for the focal phenotype. This leaves undefined the degree to which non-shared environmental influences on the focal phenotype are associated with the sorting factor (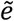in **Figure 1b**). We suggest using the term ‘idiosyncratic homogamy’. Other sources of variance can be added to the model following a similar logic, but we do not define separate terms for these. Variance attributable to correlations between social and genetic influences will intertwine the consequences of social and genetic homogamy (e.g., social homogamy can have genetic consequences)^18^, but we do not consider this its own form of homogamy.

The definitions and general framework we propose, which distinguishes the focal phenotype from its sorting factor, capture various mechanisms that were previously conceived to be distinct. For example, direct and indirect assortment are not viewed as different mechanisms. Instead, direct assortment is simply the special cases (or convenient assumption) where the focal phenotype and the sorting factor are one and the same, or more generally, where the degree of genetic, social, and idiosyncratic homogamy is equal to the relative importance of genetic, social, and idiosyncratic factors on the focal phenotype 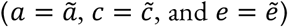. Many, but not all, prior models that assume direct assortative mating are implicitly making this assumption. It is also clear that different kinds of homogamy are not considered mutually exclusive processes but may instead be of varying importance for a given phenotype. Finally, by defining homogamy with respect to the focal phenotype, consequences of assortative mating can be derived without involving global statements about partner formation (e.g., whether assortment results from active choices or not).

### Assortative mating versus other causes of partner similarity

#### Convergence, inbreeding, and stratification

Partner similarity need not result from assortment. Because we have provided new definitions for terms related to assortative mating, it will be beneficial to also describe other causes of partner similarity, and how they relate to assortment. Note that any of the non-assortment explanations for partner similarity automatically disqualifies direct assortment as a sufficient explanation. For example, partners can become more similar over time, either because of mutual influence or because of exposure to shared environments leading to converging traits^31,41-43^ (**Figure 1c**). Convergence is not likely to be important for partner similarity in educational attainment, which is obtained relatively early in life, but may be important for other traits. If partners are similar to begin with, then some of the causes of the focal phenotype must be correlated across partners. This can come about from some form of assortative mating as described above, where covariance is induced without affecting the variance. Alternatively, it can come about through some form of shared prior cause, which will affect both variance and covariance. With a shared prior cause, the correlation between two individuals would be the same regardless of whether they were likely to mate or not. This includes inbreeding (**Figure 1d**) where some genetic variants associated with the trait are identical by decent, or its environmental equivalent: mating within an environment that also has a causal effect on their trait, but, crucially, not in a way that influences partner formation (**Figure 1e**).

Confusingly, this latter process is often described using the term ‘social homogamy’, despite it being a very different mechanism compared to assortment on social characteristics^38^. The consequences and implications differ, and confusing the two would be like confusing genetic homogamy and inbreeding. To avoid confusion, we suggest using the term ‘social stratification’ (or just ‘stratification’) to refer to environmental correlations arising from partners sharing identical causes of variance, and reserve ‘social homogamy’ for assortative mating on social characteristics, where covariance across partners arises from matching. This parallels the distinction between inbreeding (genetic variants identical by descent) and genetic homogamy (genetic variants with similar effects, but not necessarily identical). For example, if potential partners cared about the socioeconomic position of their would-be parents-in-law, then this would be social homogamy. If, on the other hand, partners tended to mate within the same city or area, and the average educational attainment happened to be different across areas, then this would be social stratification. In the first case, partner formation would be conditional on (something associated with) educational attainment, while in the latter case, partners would just happen to have similar educational attainment because they were exposed to the same cause.

This distinction can become muddled when the stratification itself is influenced by educational attainment or associated factors. Educationally segregated mating markets does not automatically imply social stratification unless these structural factors can be considered completely exogenous to the relevant traits. For example, if partner similarity in education could be wholly attributed to many would-be partners meeting at a university, then partner formation would be conditional on traits leading to university attendance, making this an example of assortative mating. Assortative mating does not by itself imply deliberate choice on behalf of the partners. Another example is non-random migration, where individuals move to areas because of their educational attainment or related factors rather than their educational attainment being decided by where they live ^44,45^. If partners mate within the same area, but the area was decided by factors related to their educational attainment, then this would constitute assortative mating per the definition above. More research is needed on distinguishing stratification and homogamy.

### Intergenerational consequences of indirect assortative mating

Under positive assortative mating, the correlation between a parent and their offspring will in most cases be inflated, regardless of whether intergenerational transmission was caused by direct influence of the parent (direct phenotypic transmission) or passively through shared genes or environments (passive genetic/environmental transmission). For example, if partners are genetically similar, then their offspring will tend to inherit alleles with similar consequences from both parents, thus inflating the genetic similarity between one parent and their offspring (and hence also phenotypic similarity)^24^. Likewise, if offspring trait values are directly influenced by their parents’ trait values, and both parents tend to have similar trait values, then the correlation between one parent and their offspring will increase because of the effect of the other parent. It is therefore important to model assortative mating when attempting to decompose the parent-offspring correlation into its various causes. Importantly, this decomposition will be sensitive to the type of assortative mating. Models assuming direct assortment, such as most extended twin models with partners, implicitly assume that all intergenerational pathways (e.g., direct transmission, passive genetic transmission, etc.,) are inflated to the same degree, proportional to the phenotypic correlation between partners. However, if, say, partners are more genetically (or environmentally) similar than implied under direct assortment, then such models may underestimate passive genetic (or environmental) transmission. Unless the degree of genetic, social, and individual homogamy happens to equal the relative importance of genetic, social, and individual factors on the focal phenotype, models assuming direct assortment may give biased and misleading results.

### Using partners of twins to model indirect assortative mating

Here, we present a method to estimate the induced correlations in causes without assuming direct assortative mating. We show how indirect assortment will affect the correlations between twins-in-law (i.e., a twin and their twin’s partner), and present partners-of-twins models that can estimate the relative importance of genetic, social, and idiosyncratic homogamy. We introduce one model that can be applied to a single generation (the iAM-ACE model), and a second model that can be applied to two generations (the iAM-COTS model). The first model can estimate the degree of genetic, social, and idiosyncratic homogamy, but cannot readily incorporate gene-environment correlations nor inform on mechanisms of intergenerational transmission. The second model allows us both to incorporate gene-environment correlations, and to investigate the consequences of indirect assortative mating on intergenerational transmission. We apply these models to educational attainment in Norwegian registry data using the extended families of 2,447 pairs of monozygotic twins, 3,360 pairs of dizygotic twins, and 206,263 pairs of full siblings, yielding 421,194 dyads of partners and in-laws. Our results indicated that partners were more genetically similar than expected under direct assortment, but that matching was primarily on sibling-shared environmental factors. When accounting for this in intergenerational models, we found that 62% of the parent-offspring correlation could be explained by passive genetic transmission, whereas the remaining was explained by direct (16%) and passive (23%) environmental transmission. Models where we assumed direct assortment fitted poorly and gave results that suggested genetic transmission accounted for almost the entire parent-offspring correlation.

## Results

### Correlations between extended family members by zygosity type

We used the Norwegian population register to identify nuclear families (children and their parents) with children born between 1975 and 1995 (and thereby old enough to have obtained education themselves). For nuclear families with more than two eligible offspring, we randomly selected two of them. We then linked nuclear families into extended family units via one of the parents’ twin or sibling, meaning an extended family unit consisted of up to eight individuals: two sets of partners in the parent generation, each with two offspring. Finally, we linked the data to administrative education registers with information on educational attainment completed by age 30. In total, we identified 212,070 extended families comprising 1,545,444individuals.

**Figure 2**and **Supplementary Table 4** shows Pearson correlations between family members stratified by zygosity group (i.e., monozygotic twins, dizygotic twins, full siblings). All family members were highly correlated for educational attainment. The correlation between monozygotic twins (.71) was higher than the correlation between dizygotic twins (.45), which in turn was higher than the correlation between full siblings (.41). Partners were somewhat more highly correlated than full siblings in all zygosity groups (∼.47). Correlations between in-laws (a twin and their twin’s partner) and co-in-laws (a twin’s partner and the other twin’s partner) were higher in monozygotic twin families than full sibling families. The parent-offspring correlation was similar in all zygosity groups (.34), as was the correlation between siblings in the offspring generation (.36). The avuncular correlation (aunt/uncle – offspring) was about equal to the parent-offspring correlation in monozygotic twin families (.34), and somewhat lower in the other family types (.23). Finally, the avuncular-in-law correlation (aunt’s/uncle’s partner – offspring) and the offspring cousin correlation was slightly higher among monozygotic twin families than other family types.

**Figure 2.**
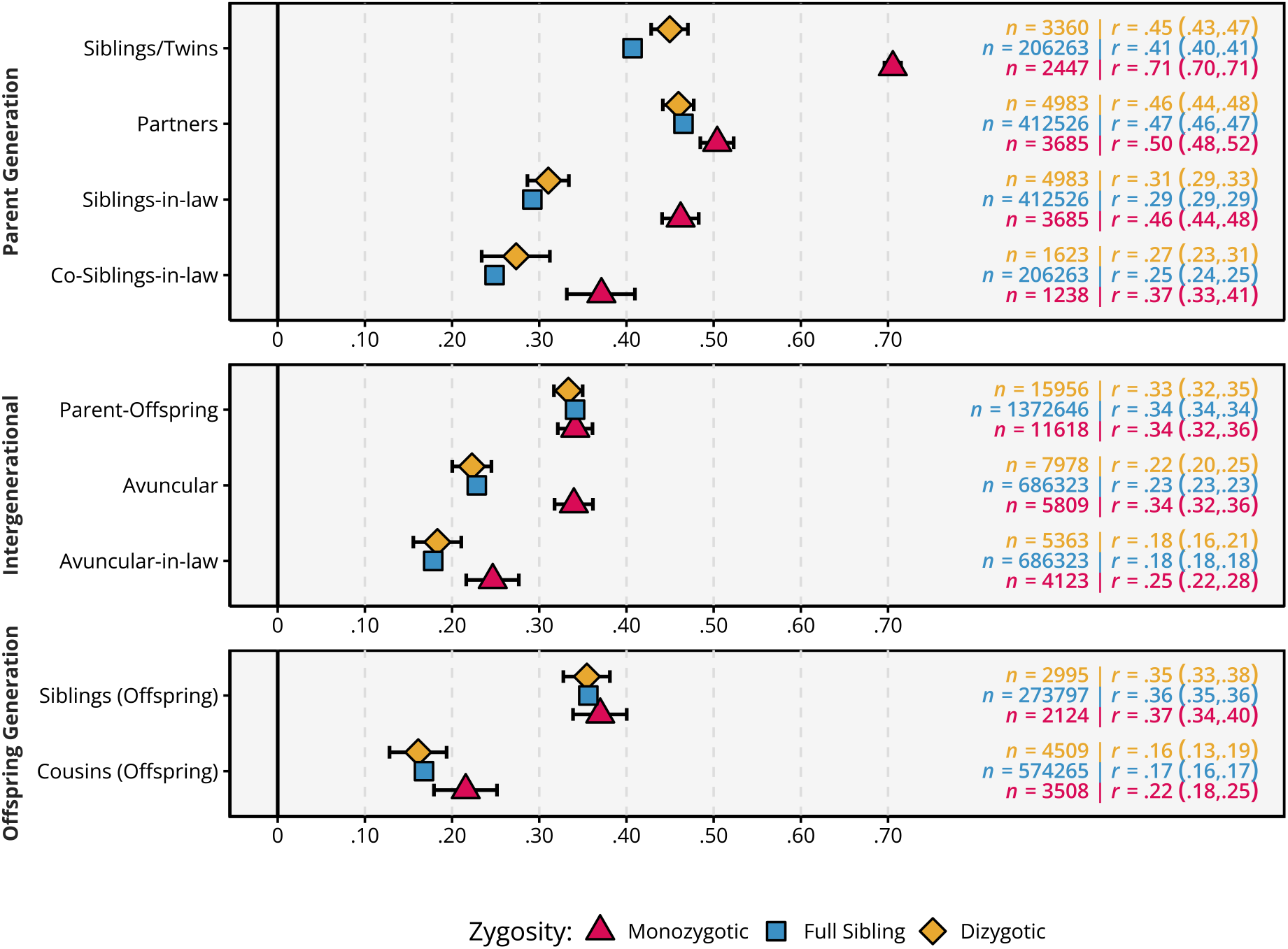
Correlations between family members’ educational attainment. (measured as years of education at age 30), stratified by zygosity type (i.e., whether siblings in the parent generation are monozygotic twins [red], dizygotic twins [yellow], or regular full siblings [blue]). Data are presented as correlations with 95% confidence intervals. Siblings-in-law = partner of sibling/twin, Co-Siblings-in-law = partner of siblings-in-law, Avuncular = Sibling/twin of parent, Avuncular-in-law = Partner of sibling/twin of parent.

### The iAM-ACE model

**Figure 3**shows the full iAM-ACE model. It is an extension of the classic twin model (called the ACE model) that includes the twins’ partners. The genetic and environmental factors (*A, C, T*, and *E*) that influence the focal phenotype with effects *a, c, t*, and *e*, respectively, are allowed to have different effects (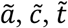, and 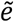, respectively) on an associated sorting factor (similar to the middle generation in the Cascade model^13^). The sorting factor is a latent variable comprising the set of traits associated with the focal phenotype that partners are assorting on. To make the model identified, one needs to constraint the variance of the sorting factor by fixing one parameter. We chose to fix ã to *a*. Note that this leads to different variance for the sorting factor and the focal phenotype, meaning that *a* and ã may still be different after standardization (i.e., after rescaling to unit variance). The assortment strength itself is indicated with the copath coefficient μ. A copath denotes similarity due to matching, such as assortative mating, and comes with special path tracing rules that allow covariance to be induced without affecting variance^28^. The degree to which the sorting factor is heritable or environmental will lead to different expected correlations between twins-in-law, which allows the model to estimate the relative importance of the different effects. For more details, see methods and **Supplementary Note 2**. We distinguish sibling-shared environmental factors (*C*, shared by both siblings and twins) from twin-shared environmental factors (*T*, shared only by twins), as earlier research has indicated that this is important for educational attainment ^35^, and the higher correlation between dizygotic twins than full siblings reported in **Figure 2** suggests the same is true in the current population. Note that sibling-and twin-shared environmental factors reflect influences that would be shared with siblings or twins, regardless of whether an individual actually has them.

**Figure 3.**
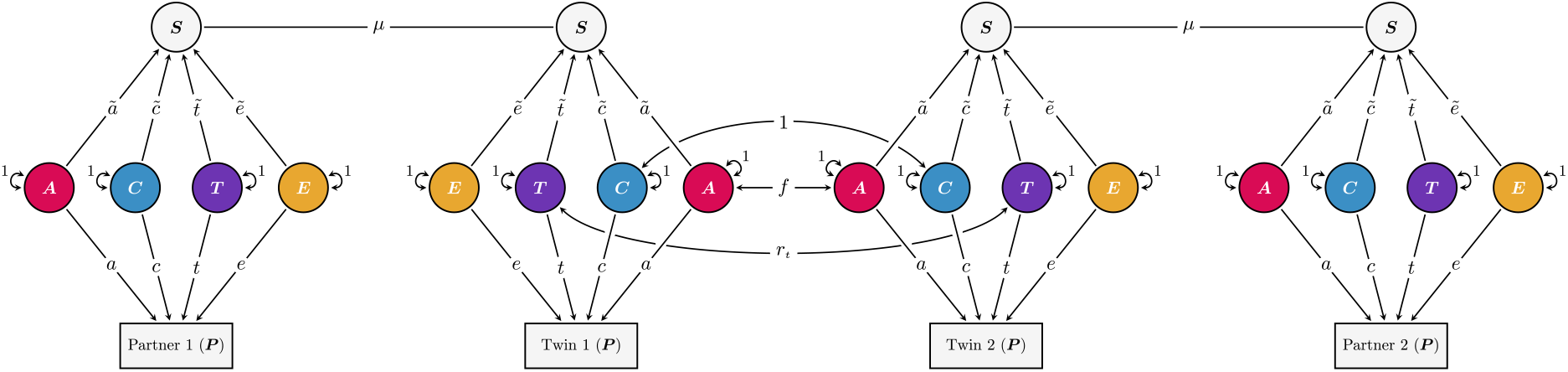
The iAM-ACE model.1. The model includes four observed individuals: a set of twins (or siblings) along with their respective partners. Differences in the observed, focal phenotype (denoted *P*) are thought to result from additive genetic factors (*A*, red), sibling-shared environmental factors (*C*, blue), twin-shared environmental factors (*T*, purple), and non-shared environmental factors (*E*, yellow). Their effects on the focal phenotype are denoted *a, c, t*, and *e*, respectively. Partners (i.e., Partner 1 – Twin 1, and Partner 2 – Twin 2) are assorting (μ) on a sorting factor (S), which are influenced by the same factors that influence the focal phenotype, albeit with different effects 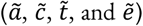. Only the relative importance of 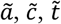, and 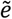 can be estimated, meaning the variance of the sorting factor must be constrained. Additive genetic factors are perfectly correlated across monozygotic twin pairs (*f*_MZ_ = 1), whereas for dizygotic twins and full siblings, the correlation is *f*_FS_ = (1 + μ ã^2^)/2 (assuming intergenerational equilibrium). The correlation in twin-shared environmental factors depend on relation (monozygotic and dizygotic twins: *r_t_* = 1, ordinary full siblings: *r_t_* = 0).

**Figure 4a** and **b** present standardized variance components (V_A_, V_C_, V_T_, and V_E_) from the iAM-ACE model. Observed educational attainment (**Figure 4a**) was estimated to be 46% (95% CIs: 41%, 50%) heritable, with significant contributions from sibling-shared (12%; 7%, 17%), twin-shared (7%; 5%, 8%), and non-shared (35%; 34%, 37%) environmental factors. The model estimated that that the observed partner correlation of .47 resulted from indirect assortment, with an estimated partner correlation of .68 (.66, .71) on the associated sorting factor. The sorting factor associated with educational attainment was, in turn, estimated to be about 38% (26%, 49%) heritable, 55% (45%, 64%) sibling-shared environment, 5% (0%, 10%) twin-shared environment, and 3% (1%, 4%) non-shared environment (**Figure 4b**). Multiplying the heritability (.38) with the estimated partner correlation (.68) gives an implied genotypic correlation between partners of .26 (.18, .34).

**Figure 4.**
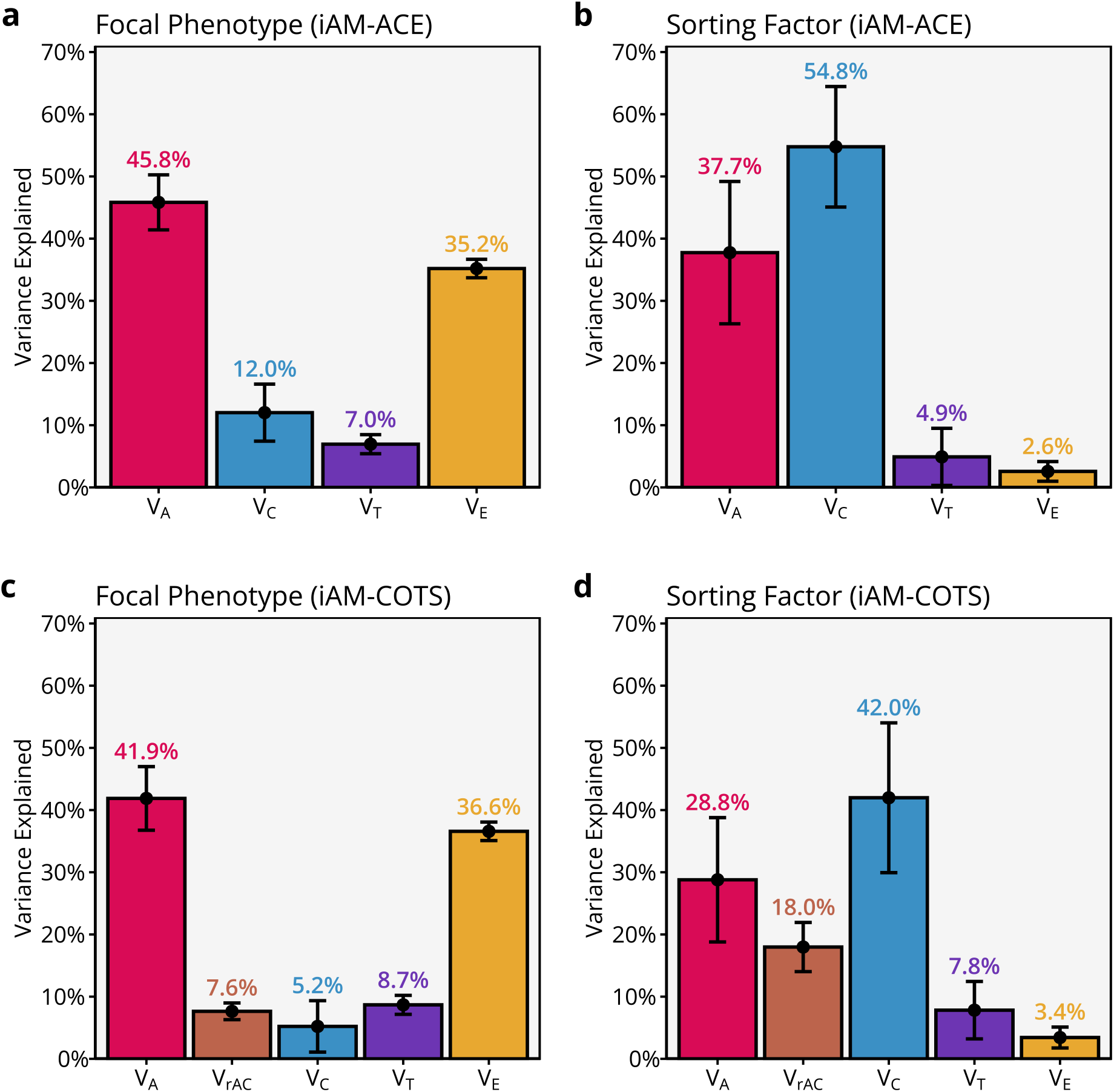
Standardized variance components for educational attainment from the iAM-ACE model (**a** and **b**) and the parent generation in the iAM-COTS model (**c** and **d**), for observed educational attainment (i.e., the focal phenotype, **a** and **c**) and the associated the sorting factor (**b** and **d**) in 212,070 extended families. Data are presented as variance components with 95% confidence intervals, where the total variance is the sum of all components. The variance components are additive genetic factors (V_A_, red), sibling-shared environmental factors (V_C_, blue), twin-shared environmental factors (V_T_, purple), non-shared environmental factors (V_E_, purple), and variance attributable to gene-environment correlations (V_rAC_, brown).

### Testing different mechanisms for partner similarity

#### Direct assortment

A model assuming direct assortment is nested within the iAM-ACE model and can therefore be tested for significant differences in fit (**Supplementary Table 5**). To assume direct assortment, the degree of genetic, social, and idiosyncratic homogamy must be constrained to equal their importance for the focal phenotype 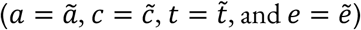, which makes the focal phenotype and the sorting factor identical. Since ã was already fixed, this removes three degrees of freedom. We find that constraining the model to direct assortment results in substantially worse fit (Δ-2LL (Δ*df* = 3) = 8598.2, *p* < .001, **Supplementary Figure 12**).

#### Direct assortment with measurement error

A special kind of indirect assortment is when the phenotype is assorted upon directly but is observed with random measurement error^46^. Because partners do not assort on measurement error (which is modelled as part of non-shared environmental influences on the observed focal phenotype), idiosyncratic homogamy will be less important than implied under direct assortment. Hence, the focal phenotype and sorting factor should have identical variance decompositions, except for non-shared environmental influences, which should be less important for the sorting factor. To test whether direct assortment with measurement error is a sufficient explanation, the model can be adapted by constraining 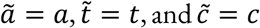 while still freely estimating 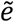. Compared with the full model, this removes two degrees of freedom. This resulted in significantly worse fit than the full model (Δ-2LL (Δ*df* = 2) = 536.9, *p* < .001).

#### Social stratification

It is possible to extend the model to include social stratification, modelled as latent environmental influences on the focal phenotype that are unrelated to partner formation and shared to the same degree by all extended family members (**Supplementary Note 3**). The model is identified because social stratification leads to smaller relative differences between the co-siblings-in-law correlation and other family members compared to expectations under assortative mating only. We find that adding social stratification does not significantly improve the fit of the model (Δ-2LL (Δ*df* = 1) < 0.001, *p* = .999).

### The iAM-COTS model

The iAM-ACE model assumes no gene-environment correlations, which, among other things, means it may underestimate the implied genotypic correlation between partners^18^. Furthermore, it is not informative about how indirect assortative mating affects intergenerational transmission. To rectify both of these shortcomings, we can extend the model. Just like the regular ACE model can be extended into a children-of-twins-and-siblings (COTS) model^33,34,47,48^, so can the iAM-ACE model be extended into the iAM-COTS model (**Figure 5**). Because children of monozygotic twins will be as related to their parent as their parent’s twin, children of twins can be used to decompose the parent-offspring correlation into parts attributable to passive genetic transmission and two forms of environmental transmission operating within and outside the nuclear family (respectively called direct phenotypic transmission and passive environmental transmission, see **Supplementary Note 4**). When both genetic and environmental transmission occur simultaneously, their effects may be correlated giving rise to gene-environment correlations. The COTS model can incorporate gene-environment correlations in the parent generation by including a covariance term, *ω*, between *A*_1_ and *C*_1_, where *ω* is constrained to equal the equivalent correlation in the offspring generation. **Figure 4c** and **d** present standardized variance components for the parent generation from the iAM-COTS model. Results are broadly comparable to results from the iAM-ACE model, with the major difference being that gene-environment correlations account for some of the variance that was previously attributed to sibling-shared environmental effects. This effect was more pronounced in the sorting factor, where gene-environment correlations were estimated to account for 18% (14%, 22%) of the variance. The remaining variance was estimated to be about 29% (19%, 39%) heritability, 42% (30%, 54%) sibling-shared environment, 8% (3%, 12%) twin-shared environment, and 3% (2%, 5%) non-shared environment. The genotypic correlation between partners was estimated to be .34 (.24, .43) in the iAM-COTS model, which is higher than in the iAM-ACE model (.26) where gene-environment correlations were missing.

**Figure 5.**
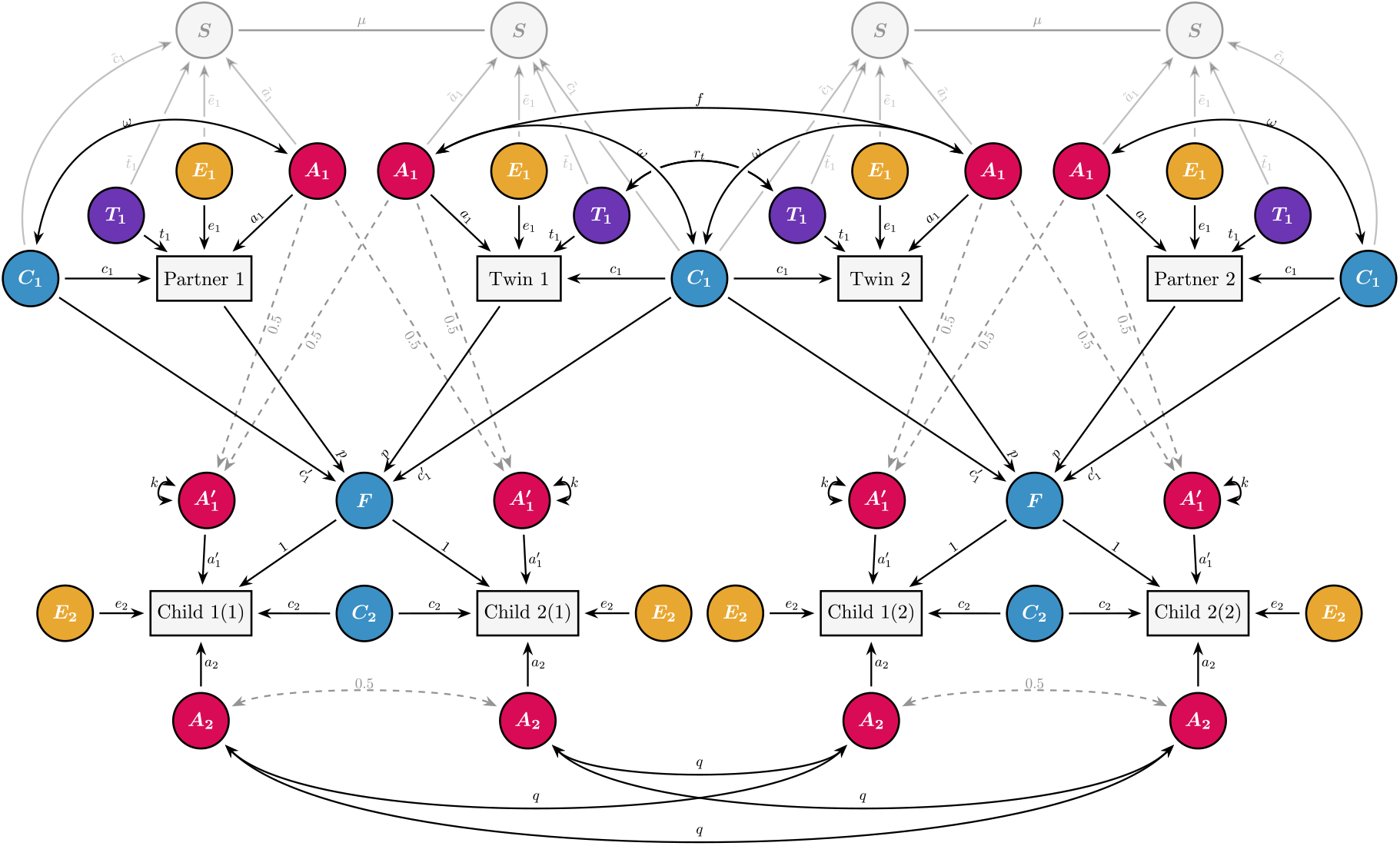
The iAM-COTS model. The model includes observations for eight individuals: a pair of twins or siblings, their respective partners, and two children each. The parent generation is equivalent to the iAM-ACE model in Figure 2 (the only difference being the inclusion of a gene-environment correlation between additive genetic effects (*A*_1_) and sibling-shared environmental effects (*C*_1_), denoted *ω*). The gene-environment correlation in the parent generation is constrained to equal that in the offspring generation: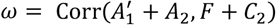. See methods and Supplementary Note 4 for more details.

### Consequences of indirect assortative mating for intergenerational modelling

**Figure 6a** presents the decomposition of the parent-offspring correlation into direct phenotypic transmission (yellow), passive genetic transmission (red), and passive environmental transmission (blue) using the iAM-COTS model. Fixing either of the three different pathways to zero resulted in significantly worse fit (**Supplementary Table 6**): no direct phenotypic transmission (Δ-2LL (Δdf = 1) = 46.4, *p* < .001), no passive genetic transmission (Δ-2LL (Δdf = 1) = 196.8, *p* < .001), and no passive environmental transmission (Δ-2LL (Δdf = 1) = 12.8, *p* < .001). The majority (.211, or 62%) of the correlation was attributed to passive genetic transmission. The second-most important pathway (.077, or 23%) was passive environmental transmission. Direct phenotypic transmission accounted for the rest of the correlation (.053, or 16%). We found that the parent-offspring correlation was substantially increased due to assortative mating, with 39% of the correlation (.133 / .341) attributable to effects via the co-parent.

**Figure 6.**
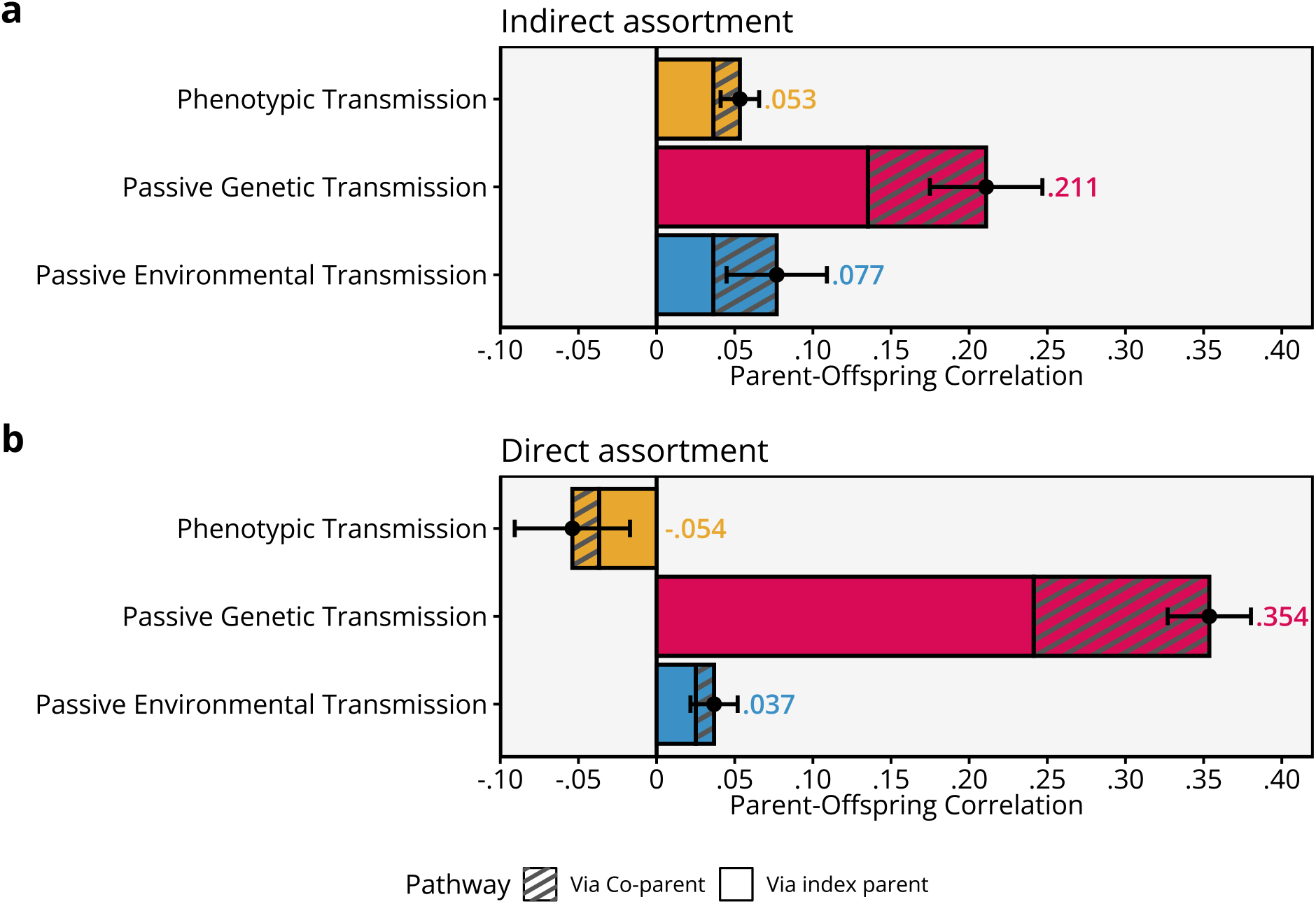
The parent-offspring correlation in educational attainment, decomposed into direct phenotypic transmission (yellow), passive genetic transmission (red), and passive environmental transmission (blue). The top panel (**a**) presents the results from the full iAM-COTS model, whereas the bottom panel (**b**) presents the results from a model constrained to assume direct assortment. Both models were applied to 212,070 extended families. Data are presented as correlation components attributable to each pathway, where the total correlation is the sum of all components. Error bars are 95% confidence intervals. The patterned areas of the bars represent the degree to which the pathway is inflated due to assortative mating.

**Figure 6b** presents a similar decomposition where the model has been constrained to assume direct assortment (the typical assumption in most COTS-models that include assortative mating). Just as in the iAM-ACE model, constraining the iAM-COTS model to direct assortment results in significantly worse fit (Δ-2LL (Δ*df* = 3) = 9299.0, *p* < .001, **Supplementary Figure 12**). Of note is that, when assuming direct assortment, genetic transmission was estimated to be substantially more important, whereas the two environmental pathways were both smaller and, in the case of direct phenotypic transmission, even reversed (resulting in a negative gene-environment correlation).

### Educational attainment in the offspring generation

**Figure 7**presents the variance decomposition of educational attainment in the offspring generation from the full iAM-COTS model. Overall, results are comparable to educational attainment in the parent generation (**Figure 4c**), with educational attainment estimated to be 41% (36%, 46%) heritable. The genetic correlation between educational attainment in the parent generation and educational attainment in the offspring generation was only estimated to .60 (.55, .64), meaning a substantial part of genetic influences on offspring educational attainment seems to be unrelated to educational attainment in the parent generation. Sibling-shared environments explained 5.1% (1.9%, 8.4%) of the variance, of which 2.2% (0.6%, 3.7%) was attributed to effects associated with parental education via passive environmental transmission, 0.4% (0.2%, 0.6%) was attributed directly to parental education via direct phenotypic transmission, 1.3% (0.7%, 2.0%) was attributed to the covariance between direct and passive environmental effects, and 1.2% (0.3%, 2.2%) was attributed to environmental effects unrelated to parental education. Finally, gene-environment correlations (7.5%; 7.1%, 7.9%) and non-shared environments (46%; 45%, 47%) accounted for the remaining variance. (Note that twin-shared environments would here form part of the non-shared environment).

**Figure 7.**
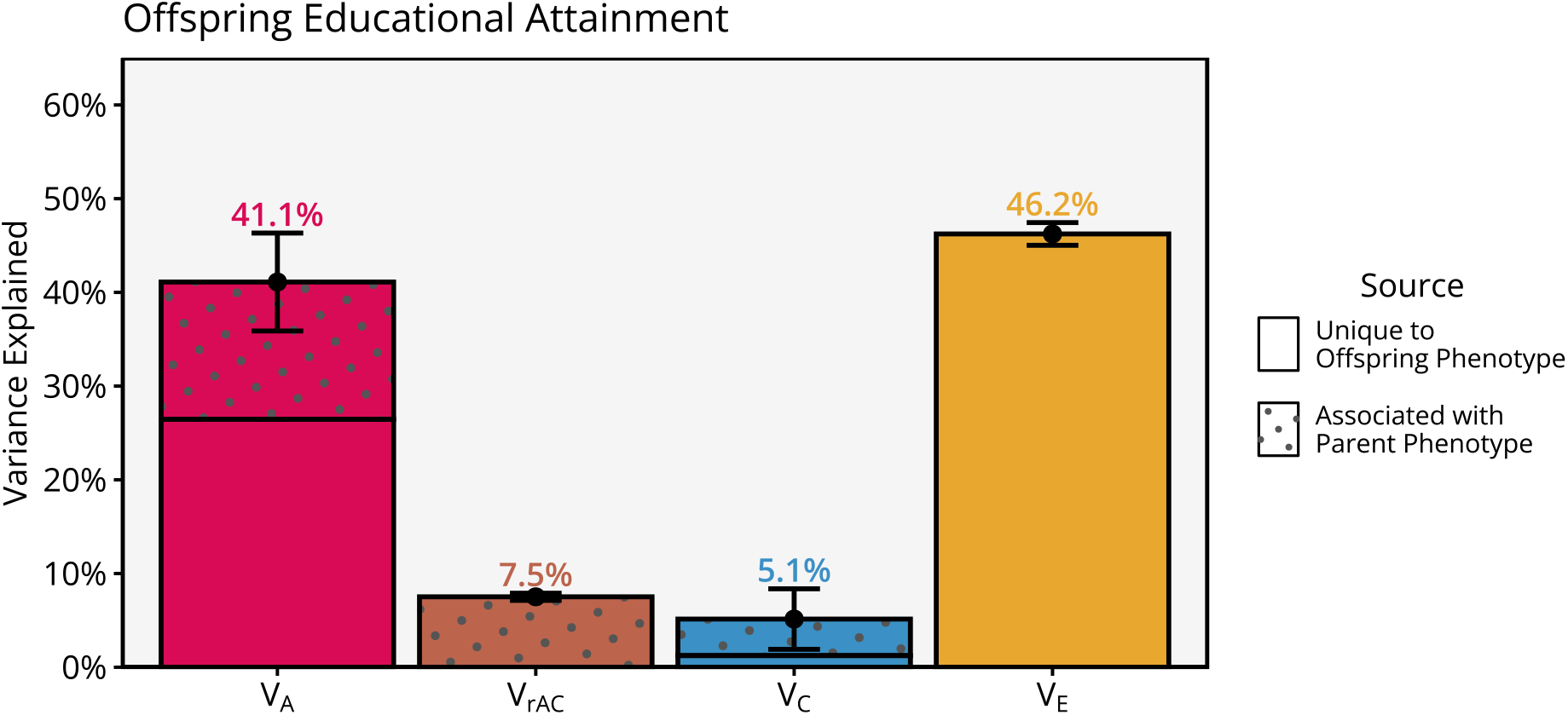
Standardized variance components for educational attainment in the offspring generation, estimated with the full iAM-COTS model applied to 212,070 extended families. Data are presented as variance components with 95% confidence intervals, where the total variance is the sum of all components. The variance components are additive genetic factors (V_A_, red), variance attributable to gene-environment correlations (V_rAC_, brown), sibling-shared environmental factors (V_C_, blue), and non-shared environmental factors (V_E_, yellow). Patterned areas of the bars indicate variance associated with parents’ educational attainment.

## Discussion

In this paper, we have developed and presented a refined framework for understanding indirect assortative mating and used it to investigate causes of partner similarity in educational attainment and its intergenerational consequences using partners of twins. In our framework, indirect assortative mating is not a distinct mechanism. Instead, it is a more general view of assortative mating where genetic and environmental components of the phenotype may or may not have different influences on partner formation than on the phenotype itself. This general framework incorporates previously distinct mechanisms, such as direct assortment, genetic homogamy, social homogamy, measurement error in the observed phenotype, and indirect assortment via associated trait(s). Overall, our empirical results replicated the finding that direct assortment is an inadequate explanation for partner similarity in educational attainment. Furthermore, our results indicated that partner similarity in both genetic and especially sibling-shared environmental factors were greater than expected under direct assortment. Finally, we have shown that attempts at decomposing intergenerational correlations are sensitive to assumptions about type of assortative mating. For traits like educational attainment, the importance of environmental transmission may be understated if failing to account for indirect assortment. When accounting for indirect assortative mating, we found that 62% of the phenotypic parent-offspring correlation could be explained by passive genetic transmission, whereas the remaining was explained by direct (16%) and passive (23%) environmental transmission.

A popular explanation for the evidence suggesting indirect assortment for educational attainment, such as higher-than-expected genetic similarity between partners, has been that it primarily results from assortment on a highly heritable, related trait like social or cognitive ability ^4,6,29^. Our results provide evidence against this hypothesis. The sorting factor was not found to be more highly heritable than educational attainment. Instead, the higher-than-expected genetic similarity between partners, which we also replicated in this analysis, could be ascribed to the high partner correlation for the sorting factor and exacerbated by gene-environment correlations. Other research also suggests that assortment on traits like cognitive performance only play a limited role in explaining partner similarity in educational attainment. For example, a recent meta-analysis found that the phenotypic correlation between partners on cognitive performance was lower than for educational attainment (.44 vs. .55), albeit with considerable between-study heterogeneity^1^. Okbay, et al. ^8^ also found that the polygenic index correlation between partners remained substantial after adjusting for cognitive performance, which implies that it is not the primary source of the increased correlation. Cognitive performance is probably measured with some measurement error, but we find it unlikely that this can fully explain the discrepancy between the hypothesis and the empirical findings.

While assortment on cognitive ability probably is an important part of the story, it does not appear to be the whole story. This is corroborated by our finding that sibling-shared environmental factors make a substantial contribution to the sorting factor: It is seemingly about five times as important for explaining partner similarity in educational attainment than for explaining educational attainment itself. This finding implies that factors shared by siblings independently of genetic similarity, such as social background or parental characteristics, is important for explaining partner similarity in educational attainment.

Non-shared environmental factors were only weakly associated with the educational sorting factor. This is not surprising considering that we found the twins-in-law correlation for monozygotic twin to be similar to the correlation between partners. Hence, we expect that obtaining higher education will only have a minor influence on partner formation, which instead seems to be determined by familial (genetic and environmental) proneness to education. This does not mean than idiosyncratic and chance events are not important for partnership formation, but that such events must be mainly unrelated to education.

Our results replicate findings from Torvik, et al. ^7^ in a different sample and with a different method. Torvik, et al. ^7^ investigated a sample of Norwegian parents of children born between 1999 and 2009, whereas we investigated Norwegian parents of children born between 1975 and 1995. Torvik, et al. ^7^ used the rGenSi model on correlations between polygenic indices and observed educational attainment among partners, siblings, and siblings-in-law, and estimated the genotypic correlation between partners to .37 (.21, .67). The estimate from the present study of .34 (.24, .43) is remarkably similar. Both the present study and Torvik, et al. ^7^ estimate the genetic similarity between partners to exceed expectations under direct assortment. (Under direct assortment, the expected genotypic correlation should be *h*^2^*μ*, the squared correlation between genotype and focal phenotype multiplied by the phenotypic partner correlation^18^, which, using the results from **Figure 4a**, should imply *r* ≈ 44% × .46 = .20). Our results are thereby broadly consistent with other studies showing greater-than-expected genetic similarity between partners, such as Robinson, et al. ^6^ and Okbay, et al. ^8^.

Our results are not consistent with Clark ^4^, who showed that, if partners were strongly assorting on a highly heritable trait associated with social class, and this resulted in a genotypic partner correlation of *r* = .57, then genetic similarity alone could explain correlations in social class among a wide range of relatives in England. First, we find the implied genotypic correlation in our study to be significantly lower than expected given his argument. Second, we find that the sorting factor was not more heritable than educational attainment itself. Instead, we found sibling-shared environmental influences to be more important, indicating considerable social homogamy. Third, our results suggest that genetic similarity could only account for 62% of the correlation between parents and offspring’s educational attainment, meaning genetic similarity alone cannot fully account for familial similarity. While there could be cultural differences between Norway and England that would change the relative importance of genetic and environmental factors, we find it unlikely that this alone can explain the discrepancy between our results and Clark ^4^. Instead, we find it probable that important mechanisms are missing from Clark’s model.

Higher social than genetic homogamy is in line with previous studies, such as Gonggrijp, et al. ^5^, Reynolds, et al. ^2^, and Collado, et al. ^3^. All three of these studies concluded that environmental partner similarity was higher than expected under direct assortment alone, and hence an important explanation for partner similarity in educational attainment. In this sense, our study can be said to replicate their results. However, this is somewhat incidental. As described in the introduction, the models used in those studies are set up in such a way where most forms of indirect assortment will look like assortment on social factors. For example, the implied genotypic correlation between partners in all these studies cannot exceed the expectation under direct assortment. The incompatibility of these studies’ results with those finding higher-than-expected genetic similarity between partners has therefore always been a foregone conclusion. The results we provide in the present study, on the other hand, are consistent with both a higher-than-expected genotypic correlation between partners and assortment on social characteristics as the most important cause of partner similarity in educational attainment.

Parents who were more highly educated had, on average, children who were more highly educated. Our results suggest that genetic similarity can account for much of this correlation (62%), but that a substantial part must be ascribed to direct (16%) and passive (23%) environmental transmission. This model provides two insights into intergenerational modelling of educational attainment. First, the decomposition of the parent-offspring correlation is sensitive to assumptions about type of assortment. Had we merely assumed direct assortment, we would have concluded that the parent-offspring correlation could be almost entirely explained by genetic transmission (**Figure 6b**). This may help explain why some earlier children-of-twins studies have reported that genetic transmission alone could account for the parent-offspring correlation, even in the same population^33^. It is important to note that the iAM-COTS model does not assume that assortment is indirect: If direct assortment was a sufficient explanation, the results would have reflected this. Our finding that a model constrained to direct assortment had poor fit to the data and gave misleading results underlines the importance of modelling assortative mating appropriately.

The second insight to note is the type of environmental transmission. Our model distinguished direct phenotypic transmission (i.e., direct effects of parental education) from passive environmental transmission (i.e., environmental factors shared by parental siblings that also influence offspring). We found that passive environmental transmission seemed more important than direct phenotypic transmission, both in terms of decomposing the parent-offspring correlation (.077 vs. .0.53) and especially in terms of variance explained in offspring educational attainment (2.2% vs. 0.3%). This suggests that important environmental effects are not necessarily stemming from the observed phenotype within the nuclear family but are instead shared by extended family members. This can include effects of grandparents, neighbourhood, or broader social background. Other studies using polygenic indices across three generations also point to extended family effects (as opposed to nuclear family effects) as a likely source of environmental effects^49^. Alternatively, the results could reflect effects of other parental phenotypes environmentally correlated with educational attainment. Future work must attempt to identify what mechanisms are involved in vertical transmission of educational attainment.

The distinction between direct and passive environmental transmission has important consequences for the expected correlations in extended families. Rao, et al. ^30^ showed how (passive) environmental and genetic transmission can be close to indistinguishable if the environmental correlation across generations is close to .50. However, genetic transmission can also be indistinguishable from direct and passive environmental transmission occurring simultaneously. Direct phenotypic transmission will typically increase the parent-offspring correlation relative to the avuncular correlation, whereas passive environmental transmission will increase the parent-offspring correlation and avuncular correlation to the same degree (thus reducing the relative difference). In an extreme scenario, they may yield a parent-offspring correlation about twice as large as the avuncular correlation, and thereby give the same expectation as a scenario where genetic transmission alone was responsible. In less extreme scenarios, they may obscure or downplay important environmental effects or otherwise give inaccurate results. Studies using phenotypic similarity between extended family members to distinguish between genetic and environmental effects may therefore give misleading results unless both direct and passive environmental transmission can be accounted for simultaneously.

A key limitation for the models presented here is that they require large amounts of data on partners of twins, even for traits with a large partner correlation like educational attainment. Furthermore, they suffer many of the same limitations as regular twin models, such as relying on the equal-environments assumption (although this is somewhat relaxed when modelling twin-shared and sibling-shared environments separately) and no gene-environment or gene-gene interactions. The iAM-COTS model can incorporate gene-environment correlations, but only by assuming it is constant across generations. Future work on the dynamics of gene-environment correlations can be beneficial^16^. For example, the consequences of and for assortative mating may depend on whether the gene-environment correlation is induced via the focal phenotype (as assumed in the iAM-COTS model), directly in the sorting factor, or via an unobserved, third variable (for example if vertical transmission is driven by an associated variable, as proposed in ^13^). Another limitation is that some individuals in the offspring generation were only observed until age 27, which may slightly underrepresent later educational attainment. Finally, like most twin studies, the models remain agnostic to what specific causal mechanisms are underlying the different variance components, leaving that for future work. On the other hand, they model all sources of variance, not merely that which is associated with observed variables such as common single nucleotide polymorphisms. Agnosticism towards mechanisms is thereby both a strength and a limitation.

In this paper, we have provided a refined framework for understanding indirect assortative mating. We do not believe this is the final say, but instead a steppingstone in the right direction. We have investigated a latent sorting factor that is fully determined by the same influences that influence educational attainment. Numerous questions remain. For example, what traits comprise the sorting factor associated with educational attainment? How relevant are preferences and active choice compared with educationally segregated mating markets^50^? What explains partner similarity in traits other than educational attainment? (Direct assortment seems to be an inadequate explanation for numerous traits^43,51^). What is the dimensionality of sorting factors across traits? Does it matter whether assortment operates primarily through matching or competition^46^? How does the sorting factor differ between men and women, and between cultures and cohorts? We believe the framework and models presented here can serve as a starting point for new, interesting research questions. We have already seen how the iAM-ACE model can be expanded to model intergenerational transmission. Other alternatives are to model other causes of partner similarity, such as convergence, which can be accomplished by including more in-laws, measured genetic data, and/or longitudinal data. Future work remains on integrating multivariate assortment and indirect assortment under a common framework, and on applying this framework to molecular genetic designs.

## Methods

### Sample

Our study is based on the Norwegian population register, which contains basic demographic information on all individuals living in Norway since 1960 (N = 8,589,458). The register includes information on, among other things, births, deaths, sex, and parentage, and can be linked to other administrative registers such as education registers (for information on educational attainment) and the Norwegian Twin Register^52^ (for information about zygosity of twin pairs). Full siblings were identified as having the same mother and father in the population register, and twins were identified as having the same mother and birth month. We only used same-sex siblings. Twins with unknown zygosity were not included as eligible siblings. Partners were defined as opposite-sex pairs who were registered as having a child together (i.e., registered as co-parents of other individuals in the population). The study was approved, and participant consent was waived by the Norwegian Regional Committee for Medical and Health Research Ethics (project #2018/ 434).

We investigated parents of children born between 1975 and 1995. Average birth year was *M* = 1956.0 (*SD* = 7.3) for fathers and *M* = 1958.7 (*SD* = 6.9) for mothers. We limited our analyses to Norwegian-born families with available educational data on both parents and children. We identified nuclear family units via shared parentage, randomly choosing two children for larger nuclear families. We then linked the nuclear families into extended family units via one of the parents’ twin or sibling. We first linked together units by monozygotic and dizygotic twins and included twin uncles and aunts without eligible children themselves (to increase statistical power). For the remaining nuclear families, we linked them together via one of the parents’ siblings, choosing randomly if there were multiple candidates. No individual formed part of more than one extended family unit. This procedure resulted in 212,070 extended families, of which 2,447 included monozygotic twins, 3,360 included dizygotic twins, and the remaining 206,263 included full siblings. In total, our analysis comprised 1,545,444 unique individuals: 845,334 (50% women) in the parent generation and 700,110 (49% women) in the offspring generation. Sample sizes for each dyad within each zygosity group are presented in **Figure 2**. Sample sizes for each cell in the covariance matrices are presented in **Supplementary Figure 11**.

### Measures

Individual-level data on educational attainment was provided by Statistics Norway and was available yearly from 1980 up to and including 2021. Educational attainment was recoded into years of education and then z-standardized within sex and generation. For each individual, we used the highest attained educational attainment recorded by the age of 30. For individuals with no records before that age (e.g., already older than 30 in 1980), we used the earliest available record. We had access to some data from before 1980, although this was incomplete and considered by some to be unreliable. We only used this data for individuals with no data after 1980. In those cases, we used the latest entry. For individuals born between 1992 and 1995 (who had not reached the age of 30 by the end of 2021), we used the latest available entry.

### Analyses

We estimated Pearson correlations between all unique family relationships separately within each zygosity group. To account for dependencies (some individuals forming part of multiple dyads), we used maximum likelihood to estimate a semi-constrained covariance matrix in OpenMx^53^ 2.20.6, using R^54^ 4.2.3, and standardizing to correlations afterwards. Equivalent relations (e.g., Sibling 1 – Partner 2 and Partner 1 – Sibling 2) were constrained to be equal, as was means and variances within generations. We estimated 95% confidence intervals using Fisher’s *r*-to-*z* transformation, applying the Wald method (± *z_α/2_*× *SE* in the *z* scale) and back-transforming to the correlation scale. Correlations are reported in **Figure 2** and **Supplementary Table 4**. For unconstrained correlation matrices, see **Supplementary Figure 11**. The iAM-ACE model and iAM-COTS model (described below) were also estimated on the raw data using OpenMx^53^ 2.20.6, using R^54^ 4.2.3. For model estimates, we estimated 95% confidence intervals using the Wald method (± *z_α/2_*× *SE*). Nested models were compared using log-likelihood ratio tests. The scripts containing the models, together with result files and scripts for reproducing the figures, are available at https://osf.io/dznbk/.

#### The iAM-ACE model

The iAM-ACE model, illustrated in **Figure 3**, is a structural equation model that uses observed covariances between partners, twins, twins-in-law, and co-twins-in-law across zygosity groups (monozygotic twins, dizygotic twins, and full siblings) to differentiate the relative importance of genetic and environmental factors on the observed phenotype from their relative importance for assortative mating. The model is described in more detail in **Supplementary Notes 1 and 2**. Differences in the observed, focal phenotype (denoted *P*) are thought to result from additive genetic factors (*A*), sibling-shared environmental factors (*C*), twin-shared environmental factors (*T*), and non-shared environmental factors (*E*). Their effects on the focal phenotype are denoted *a, c, t*, and *e*, respectively. Partners (i.e., Partner 1 – Twin 1, and Partner 2 – Twin 2) are assorting (*μ*) on a latent sorting factor (S), which are influenced by the same factors that influence the focal phenotype, albeit with different effects 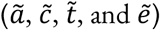. Only the relative importance of 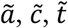 and 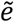 can be estimated, meaning the variance of the sorting factor must be constrained. We accomplished this by fixing ã to equal *a* and rescaling the parameters afterwards. Additive genetic factors are perfectly correlated across monozygotic twin pairs (*f*_MZ_= 1), whereas for dizygotic twins and full siblings, the correlation is *f*_FS_= (1 + *μ* ã ^2^)/2(assuming intergenerational equilibrium). The correlation in twin-shared environmental factors depend on relation (monozygotic and dizygotic twins: *r_t_*= 1, ordinary full siblings: *r_t_*= 0). Expected covariances are listed in **Supplementary Table 1** for path specification and **Supplementary Table 2** for variance specification. We used the path specification when applying the model to data. Possible extensions to the models (cross-trait assortment and social stratification) are described in **Supplementary Note 3**. Simulations of the model are presented in **Supplementary Note 5**. Correlations between parameter estimates are presented in **Supplementary Figure 13**.

#### The iAM-COTS model

The iAM-COTS model, illustrated in **Figure 5**, is an extension of the iAM-ACE model where two children per partnership have been added to the model. Expected covariances are listed in **Supplementary Table 3**. The parent generation is equivalent to the iAM-ACE model in Figure 2(the only difference being the inclusion of a gene-environment correlation, denoted *ω*). To differentiate factors that operate on the different generations, each variable and its effect have been subscripted with 1 if they operate in the parent generation (i.e., if they were included in the iAM-ACE model) and 2 if they operate exclusively in the offspring generation. The offspring phenotype is decomposed similarly to the parental phenotype, albeit with no twin-shared environments (i.e., with additive genetic, sibling-shared environmental, and non-shared environmental factors). The additive genetic factor is split into the component that is associated with the parental phenotype 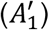 and the component that is unique to the offspring phenotype (*A*_2_). The offspring genetic factor associated with the parental phenotype is a function of the parental genetic factors and recombination variance (denoted *k*, equal to 1 − *f*_FS_in intergenerational equilibrium). The other genetic factor, *A*_2_, is correlated between siblings (.50) and cousins (*q*). The genotypic correlation between cousins will depend on whether their parents are monozygotic twins or not (*q*_MZ_= .25; *q*_DZ_= *q*_FS_= .125).

The sibling-shared environmental influences are also split into that which is associated with the parental phenotype via some form of environmental transmission (*F*) and that which is unique to the offspring generation (*C*_2_). The model includes two forms of environmental transmission: direct phenotypic transmission where the parental phenotype influences the offspring phenotype directly (*P*), and passive environmental transmission where the sibling-shared environmental factor that influenced the parents also influence the offspring 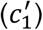. If both genetic and environmental transmission are non-zero, the effects of 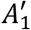 and *F* can become correlated (i.e., a gene-environment correlation). The model uses this gene-environment correlation as a best guess for what the gene-environment correlation is in the parent-generation, such that the correlation between *C*_1_ and *A*_1_, denoted *ω*, is constrained to equal the correlation between 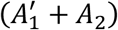 and (*F* + *C*_2_).

The key correlation of interest is that between parents and offspring. In the iAM-COTS model, it is modelled as the sum of components attributable to passive genetic transmission, passive environmental transmission, and direct phenotypic transmission. In **Supplementary Note 4**, we describe the equation that represents the parent-offspring covariance and show how the avuncular covariance across zygosity groups can be used to estimate the various components. Simulations of the model are presented in **Supplementary Note 5**. Correlations between parameter estimates are presented in **Supplementary Figure 13**.

## Data Availability

The raw data are protected and are not available due to data privacy laws. The data for this study encompasses educational outcomes and demographic information for entire cohorts of the Norwegian population. Researchers can access the data by application to the Norwegian Regional Committees for Medical and Health Research Ethics and the data owners (Statistics Norway and The Norwegian Institute of Public Health). The authors cannot share these data with other researchers.

## Code Availability

Code is available at https://osf.io/dznbk/.

## Acknowledgements

This work is part of the REMENTA and PARMENT projects and was supported by the Research Council of Norway (#300668 and #334093, respectively, to F.A.T.). This work was partly supported by the Research Council of Norway through its Centres of Excellence funding scheme (#262700). Data on twin zygosity were obtained from the Norwegian Twin Registry, Norwegian Institute of Public Health. This work is a part of the European Research Council (ERC) consolidator grant GeoGen “The PsychoGeography of Intergenerational Mobility: Early life socioeconomic position, mental health, and educational performance” (Grant agreement No. #101045526). This work was co-funded by the European Union (ERC, BIOSFER, 101071773). Views and opinions expressed are however those of the author(s) only and do not necessarily reflect those of the European Union or the European Research Council. Neither the European Union nor the granting authority can be held responsible for them. This work was performed on the TSD (Tjenester for Sensitive Data) facilities, owned by the University of Oslo, operated and developed by the TSD service group at the University of Oslo, IT-Department (USIT). We thank M. Keller, T. Chen and G. Bignardi for their helpful comments.

## Author Contributions

The authors jointly conceived of the research question. H.F.S. designed the models, carried out the analyses, and wrote the manuscript, including the supplementary information. E.M.E and F.A.T supervised the project, provided critical feedback, discussed the results, and helped shape the manuscript.

