## Supplementary Information for "Understanding indirect assortative mating and its intergenerational consequences for educational attainment"

Hans Fredrik Sunde[1*], Espen Moen Eilertsen[2], Fartein Ask Torvik[1,2]

[1] Centre for Fertility and Health, Norwegian Institute of Public Health, Oslo, Norway
[2] PROMENTA Research Centre, Department of Psychology, University of Oslo, Oslo, Norway

### Supplementary Note 1: Direct and indirect assortative mating

#### Introduction to path tracing rules

Throughout this paper, we will employ path diagrams to understand the consequences of varying causes of partner similarity. Path diagrams consist of nodes and edges[1-3]. Nodes represent variables and are indicated with squares for observed variables and circles for unobserved variables. Edges represent the causal links between the different variables, where single-headed arrows represent causation in the direction of the arrows (i.e., X → Y means that X causes Y) and double-headed arrows represent exogenous sources of variation and covariation (i.e., X ↔ Y means that something outside the model causes covariance between X and Y). Edges have path coefficients, which denote the strength of the relationship between the two variables attributable to that which is represented by that edge. The expected covariance between any two variables can be found by applying path tracing rules. These involve tracing each valid chain of paths between the two variables, multiplying the path coefficients in each chain, and summing together all valid chains. A valid chain always starts tracing up against the direction of arrows (←) and includes exactly one double-headed arrow (↔). After tracing through a double-headed arrow, the chain changes direction and you must trace with the direction of arrows (→). A chain can only change direction once (using a double-headed arrow), meaning you are not allowed to trace forwards and then backwards. The variance of a variable can be considered its covariance with itself and can be found using the same rules.

A third type of edge is the co-path, which denotes covariance attributable to matching (e.g., assortative mating) where covariance is induced without causing variance[4-6]. Such matching will induce correlations in all the causes of the variables that are matched (along with all other associated variables). To account for this, the co-path comes with special path tracing rules: A co-path connects two valid chains per standard path tracing rules into longer chains (↔ — ↔). A chain cannot start or end with a co-path (you must always start tracing backwards against an arrow, ←), and a given co-path can only be included once in each chain. We will demonstrate the path tracing rules below.

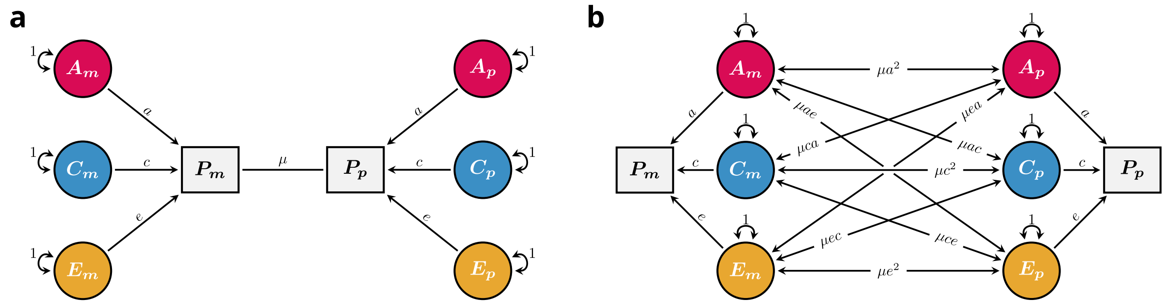

**Supplementary Figure 1:** Assortative mating leads to correlation in causes. **a**) Direct assortative mating illustrated with a copath, denoted $\mu$, connecting the two focal phenotypes, denoted $P$. The phenotype is assumed to be influenced by additive genetic, sibling-shared environmental and non-shared environmental influences, denoted $A$, $C$, and $E$, respectively. **b**) Resulting correlations in causes by direct assortment on $P$.

#### Direct assortative mating

**Supplementary Figure 1a** shows a path diagram with two partners ($m$ and $p$) whose observed phenotypes ($P$) are caused by three uncorrelated factors: additive genetic factors ($A$), sibling-shared environmental factors ($C$), and non-shared environmental factors ($E$). The importance of the different causes is denoted with path coefficients $a$, $c$, and $e$, respectively, just like classical twin models[7]. While more complicated models are possible, for example with gene-environment correlations or sex-specific effects, we will use a simple model for now to ease the explanation. To find the phenotypic variance, $Var(P)$, we can trace all valid chains leading from, say, $P_m$, and back to itself. There are three such chains ($a \times 1 \times a$, $c \times 1 \times c$, and $e \times 1 \times e$), meaning the phenotypic variance can be expressed as $\sigma_P^2 = a^2 + c^2 + e^2$. (From here on, we will omit the unit path coefficients, and, for the most part, skip directly to the simplified expressions).

We suppose that partners are assorting directly on $P$, indicated with the co-path ($\mu$) between $P_m$ and $P_p$. To find the covariance between $P_m$ and $P_p$, we must trace all valid chains between them. Because this will involve all chains going from $P_m$ (and $P_p$) back to itself, we can employ a shortcut where we bundle this set of chains together before traversing the co-path. To find the covariance, we can therefore first trace all valid paths going from $P_m$ and back to itself (i.e., its variance, $\sigma_P^2$), multiply them with the co-path coefficient ($\mu$), and finally multiply the result with all valid paths going from $P_p$ and back to itself (i.e., its variance, $\sigma_P^2$). The phenotypic covariance between the two partners can therefore be expressed as:

$$Cov(P_m, P_p) = \mu(a^2 + c^2 + e^2)^2 = \mu\sigma_P^4 \qquad (1)$$

If we imagine that all variables have unit variances (i.e., $\sigma_P^2 = 1$), which we will do throughout this paper, covariances will be equal to correlations and can be interpreted as such. In this scenario, **Equation 1** happens to simplify to $\mu$, meaning $\mu$ will equal the phenotypic correlation between partners ($\rho_{partner}$). However, to understand the implications of assortative mating, it is important to not confuse the co-path coefficient with the phenotypic partner correlation. If we instead expand **Equation 1**, we see that the phenotypic correlation can be expressed as the sum of the induced correlations in causes (**blue**), weighted by the importance of those causes (**red**):

$$\rho_{partner} = a^2(\mu a^2) + c^2(\mu c^2) + e^2(\mu e^2) + ac(2a\mu c) + ae(2a\mu e) + ce(2c\mu e) \qquad (2)$$

Assortment on $P$ will lead to partner correlations in all the causes of $P$, as illustrated in **Supplementary Figure 1b**. For example, if the phenotype is heritable, then the genetic factors should be correlated across partners ($Corr(A_m, A_p) = \mu a^2$). Numerous studies have documented genetic similarity between partners at loci associated with educational attainment, which is taken as evidence of assortative mating[8-12]. Genetic similarity between partners will, among other things, increase genetic similarity between relatives (including dizygotic twins) in subsequent generations, thereby biasing twin models unless accounted for. Similar biases exist in other type of genetic models. Attempts to account for assortative mating in such models typically assume direct assortment, for example by using the implied genetic similarity between partners and adjusting the model accordingly.

#### Indirect assortative mating

The key assumption under direct assortment is that the induced correlations in causes will be proportional to the importance of those causes. This is evident in **Equation 2**, where the terms in the induced correlations in causes (**blue**) are the same as the terms denoting the importance of those causes (**red**). A more general framework for assortative mating would relax this assumption. To accomplish this, we can adapt the model in **Supplementary Figure 1a** by distinguishing the observed phenotype in question (denoted $P$, henceforth called *the focal phenotype*) from its *sorting factor* (denoted $S$). **Supplementary Figure 2** illustrates such a model. Here, the sorting factor is influenced by the same factors that influence the focal phenotype. This is similar to the Cascade model in Keller, et al. [6], and allows us to conceptually distinguish the importance of different causes for the phenotype (denoted $a$, $c$, and $e$) from their importance for assortative mating (denoted $\tilde{a}$, $\tilde{c}$, and $\tilde{e}$). We will provide examples later.

The sorting factor can be seen as representing the associated trait or set of traits undergoing assortative mating, as they relate to the focal phenotype. Residual genetic and environmental influences on the associated traits that are not associated with the focal phenotype is, by definition, not relevant for partner similarity in the focal phenotype, and is therefore not included in the sorting factor (but see **Supplementary Note 3**). That is, the genetic and environmental correlations between the focal phenotype and its sorting factor are always 1, meaning the sorting factor only includes the components of the associated traits that are associated with the focal phenotype.

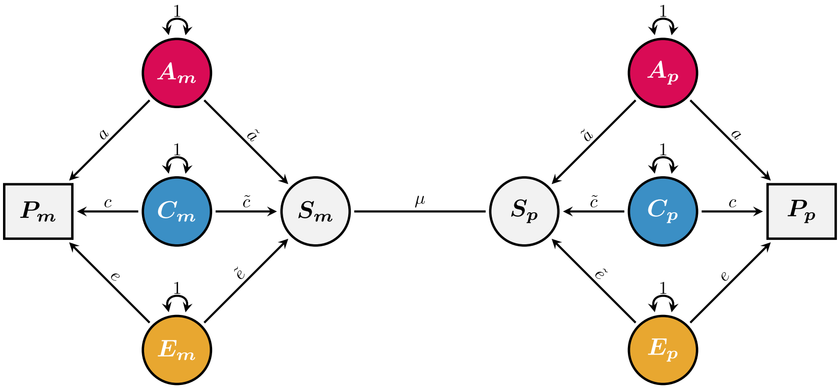

**Supplementary Figure 2:** Partner correlations in the focal phenotype ($P$) resulting from assortment on an associated sorting factor ($S$), which are caused by the same genetic and environmental factors as the focal phenotype.

Using path tracing rules, we can express the variance of the sorting factor as $\sigma_S^2 = \tilde{a}^2 + \tilde{c}^2 + \tilde{e}^2$. The consequences of assortative mating for the focal phenotype depend on the strength of assortment on the sorting factor and the nature of the correlation between the focal phenotype with its sorting factor. In **Supplementary Figure 2**, there are three pathways that contribute to this correlation, each corresponding to the three sources of variance:

$$Corr(P, S) = a\tilde{a} + c\tilde{c} + e\tilde{e} = \sigma_{PS} \quad (3)$$

(Again, it is possible to include other sources of covariance, such as gene-environment correlations, but we have omitted that here to ease the explanation). If partners are assorting directly on the focal phenotype, then $a = \tilde{a}$, $c = \tilde{c}$, and $e = \tilde{e}$, which in turn means that $\sigma_P^2 = \sigma_{PS} = \sigma_S^2$. In other words, drawing the assortment process as in **Supplementary Figure 2** does not preclude direct assortative mating. Instead, **Supplementary Figure 2** presents a more general framework for understanding assortative mating, of which direct assortment is merely a special case. We will present other hypothetical cases below.

In **Supplementary Figure 2**, $\mu$ denotes the strength of assortment on the partners' sorting factors ($S_m$ and $S_p$). Using path tracing, we find that the partner covariance between the sorting factors is $Cov(S_m, S_p) = \mu\sigma_S^4$. When we are assuming unit variances ($\sigma_S^2 = 1$), $\mu$ will equal the partner correlation on the sorting factor. If we shift our focus back to the focal phenotype, we find that the correlation between partners is the strength of assortment multiplied by the squared correlation between the sorting factor and the focal phenotype:

$$\rho_{partner} = \mu(a\tilde{a} + c\tilde{c} + e\tilde{e})^2 = \mu\sigma_{PS}^2 \qquad (4)$$

Note that the partner correlation on the sorting factor will always be equal or larger than on the focal phenotype ($|\mu\sigma_S^4| \geq |\mu\sigma_{PS}^2|$). Just as we did earlier, we can expand **Equation 4** to express the phenotypic partner correlation as the sum of the induced correlations in causes (**blue**), weighted by the importance of those causes (**red**):

$$\rho_{partner} = a^2(\mu\tilde{a}^2) + c^2(\mu\tilde{c}^2) + e^2(\mu\tilde{e}^2) + ac(2\tilde{a}\mu\tilde{c}) + ae(2\tilde{a}\mu\tilde{e}) + ce(2\tilde{c}\mu\tilde{e}) \qquad (5)$$

The difference in **Equation 5** compared to **Equation 2** is that the induced correlations in causes are no longer necessarily proportional to the causes' association with the focal phenotype. Instead, the relevant parameters are their association with the sorting factor.

##### Hypothetical examples of indirect assortment.

Let us consider a few situations where the consequences differ from direct assortment.

First, consider a situation where the phenotype is assorted upon directly, but is observed with measurement error[13]. This can be either because of misclassification or merely a crude measurement instrument such as a Likert scale. (Educational attainment can be likened to a Likert scale with only a few response options and will have measurement error if it is interpreted as a measure of, say, educational potential, even if everyone is classified correctly). In such a scenario, the unique environmental factor ($E$) in **Supplementary Figure 2** will consist of true environmental influences and noise. Partner formation is presumably random with respect to noise, meaning the cross-partner correlation between the unique environmental factors will be less than implied by **Supplementary Figure 1b** (i.e., $Corr(E_m, E_p) < \mu e^2$). Because the phenotypic partner correlation is the weighted sum of the induced correlations in causes (**Equation 2 and 5**), the other correlations in causes must be higher than implied by **Supplementary Figure 1b** to compensate. Consequently, models of direct assortment assume that there is no measurement error in the measured phenotype[13], and assuming direct assortment when there is measurement error may bias results by, for example, underestimating genetic similarity between partners.

In the above situation, the sorting factor would simply be the focal phenotype without measurement error. Let us entertain another situation where partners were assorting solely on sibling-shared environmental factors, such as social background, and that other influences on the focal phenotype are incidental with respect to partner formation. This scenario is sometimes called (pure) social homogamy. In this hypothetical situation, the sorting factor is social background (or to be more precise, the aspects of social background that are associated with the focal phenotype), and $\tilde{c} = 1$, $\tilde{a} = 0$, and $\tilde{e} = 0$. The phenotypic partner correlation would be solely attributable to the induced correlation in sibling-shared environmental factors, $\rho_{partner} = c^2(\mu\tilde{c}^2)$, meaning the true shared-environmental similarity must be much greater than implied by **Equation 2**. Likewise, the genetic similarity would be zero, despite **Equation 2** implying otherwise.

A third and final hypothetical example is close to what is sometimes called (pure) genetic homogamy, where environmental influences are incidental with respect to partner formation. Suppose for the sake of argument that the focal phenotype (educational attainment) was fully determined by cognitive ability, social background, and randomness. Suppose further that educational attainment and cognitive ability were correlated exclusively because of a genetic correlation (meaning environmental influences on cognitive ability was not relevant for educational attainment). Finally, suppose that partners were solely assorting on cognitive ability. In this scenario, the sorting factor is cognitive ability (or to be more precise, the aspects of cognitive ability that are associated with educational attainment), and the sorting factor and focal phenotype are correlated exclusively through genetic pathways, meaning $\tilde{a} = 1$, $\tilde{c} = 0$, and $\tilde{e} = 0$. Here, the phenotypic partner correlation would be solely attributable to the induced correlation in genetic factors, $\rho_{partner} = a^2(\mu\tilde{a}^2)$, meaning the true genetic similarity must be much greater than implied by **Equation 2**.

The consequences of (pure) genetic homogamy as in the third example were laid out by both Fisher[14] and Wright[3,15]. They also laid out the consequences for direct assortment, but neither considered cases where environmental influences were only partly incidental. That is, they only considered cases where the implicit assumption was either $a = \tilde{a}$ or $\tilde{a} = 1$. However, in many if not most cases, neither direct assortment, pure genetic homogamy, nor pure social homogamy are likely to be adequate descriptions. Even if multiple phenotypes are observed, the true sorting factor(s) may be unobserved, meaning $\tilde{a}$, $\tilde{c}$, and $\tilde{e}$ will still be unknown unless strong assumptions are imposed on the research design.

#### Consequences for twins-in-law

In the preceding pages, we have separated the induced correlation in causes (e.g., $\mu\tilde{a}^2$) from the importance of those causes for the phenotype (e.g., $a^2$). The next task is to find a way to estimate the induced correlation in causes without having to assume direct assortative mating. There may be numerous ways to accomplish this, depending on the research design and specific operationalisation of genetic, social, and idiosyncratic homogamy. Here, we show how twins-in-law can inform on this issue.

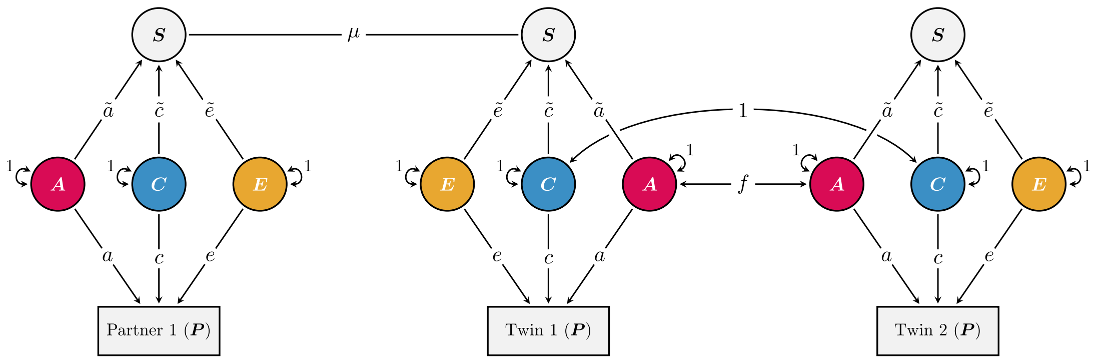

**Supplementary Figure 3** Extending the path diagram in Supplementary Figure 2 with the twin of one of the partners. Subscripts have been removed to reduce clutter. This path diagram can be estimated in a structural equation model provided (1) that the variance of S is constrained, and (2) groups with different genotypic correlations ($f$) are included (e.g., monozygotic and dizygotic twins).

**Supplementary Figure 3** extends **Supplementary Figure 2** by including one of the partners' twin (or full sibling). Sibling-shared environmental factors ($C$) are, by definition, perfectly correlated between twins, whereas the genotypic correlation (denoted $f$) depends on zygosity. In the classic twin model (henceforth, *the ACE model*), this would be $f_{MZ} = 1$ for monozygotic twins and $f_{DZ} = 1/2$ for dizygotic twins and full siblings. Assortative mating in earlier generations will lead to increased genotypic correlation between dizygotic twins and full siblings, which can be accounted for by adjusting $f_{DZ}$ according to the implied genotypic correlation between partners. However, to simplify the following explanation, we will assume $f_{DZ} = 1/2$ for now. Using path tracing rules, we find that the phenotypic correlation between twins can be expressed as:

$$\rho_{twin} = fa^2 + c^2 \tag{6}$$

This is just like in any other classical twin model. Where it gets interesting is the similarity between twins-in-laws, and by extension, other family members: Using path tracing, we find that the correlation between twins-in-law (i.e., Partner 1 ↔ Twin 2) can be expressed as:

$$\begin{aligned} \rho_{inlaw} &= (a\tilde{a} + c\tilde{c} + e\tilde{e})\mu(af\tilde{a} + c\tilde{c}) \\ &= \mu\sigma_{PS}(af\tilde{a} + c\tilde{c}) \end{aligned} \tag{7}$$

Notably, this expression includes the induced correlation in genetic and environmental factors ($\tilde{a}$ and $\tilde{c}$). Correlations between monozygotic and dizygotic twins contain enough information to calculate the relative importance of $a$, $c$, and $e$. Correlations between monozygotic and dizygotic twins-in-law add two new degrees of freedom, which can be used to estimate the relative importance of $\tilde{a}$, $\tilde{c}$, and $\tilde{e}$. Because the sorting factors are not observed, there is not enough information to calculate their absolute values, but we can find their relative importance if we, for example, assume the sorting factor has unit variance (meaning $\tilde{e} = \sqrt{1 - \tilde{a}^2 - \tilde{c}^2}$). This is not necessary for the focal phenotype, where the observed variance provides the necessary information to estimate the absolute values of both $a$, $c$, and $e$.

Just as it is possible to use Falconer's equations to quickly calculate the heritability of the focal phenotype, it is possible to perform a back-of-the-envelope calculation to find the relative importance of the three pathways between the focal phenotype and sorting factor. This can be achieved by using the sibling-in-law correlations for monozygotic and dizygotic twins in conjunction with the partner correlation in the following equations:

$$\frac{a\tilde{a}}{\sigma_{PS}} = \frac{2(\rho_{inlaw(MZ)} - \rho_{inlaw(DZ)})}{\rho_{partner}} \qquad (8a)$$

$$\frac{c\tilde{c}}{\sigma_{PS}} = \frac{2\rho_{inlaw(DZ)} - \rho_{inlaw(MZ)}}{\rho_{partner}} \qquad (8b)$$

$$\frac{e\tilde{e}}{\sigma_{PS}} = \frac{\rho_{partner} - \rho_{inlaw(MZ)}}{\rho_{partner}} \qquad (8c)$$

where $\rho_{inlaw(MZ)}$ and $\rho_{inlaw(DZ)}$ denote the sibling-in-law correlations for monozygotic and dizygotic twins, respectively, and $\rho_{partner}$ denotes the partner correlation. Just like Falconer's equations, these are somewhat biased when assortment has been ongoing in previous generations (i.e., if $f_{DZ} > 1/2$), but still give some useful intuitions: If $\rho_{partner}$ is larger than $\rho_{inlaw(MZ)}$, then the sorting factor must be partly attributable to unique environmental influences. If the $\rho_{inlaw(MZ)}$ is larger than $\rho_{inlaw(DZ)}$, then the sorting factor must be partly attributable to genetic influences. And if $\rho_{inlaw(DZ)}$ is more than half of $\rho_{inlaw(MZ)}$, then – assuming random mating in preceding generations – the sorting factor must be partly attributable to shared environmental influences. Finally, under direct assortment where $a = \tilde{a}$ and $c = \tilde{c}$, the twins-in-law correlation will be the product of the partner correlation and twin correlation. Incorporating $f_{DZ} \neq 1/2$ is straightforward in a structural equation model, as is expanding the model to include, say, gene-environment correlation. However, the intuitions one can glean from looking at the correlations alone are less straightforward.

The conceptualisation presented here, where the focal phenotype is separated from its sorting factor, is similar to the Cascade model presented in Keller, et al. [6]. However, they initially reported that there was not enough information to estimate the parameters going to the sorting factor. Instead, they suggested to either fix the parameters to equal the respective parameters for the focal phenotype, or to fix different parameters to zero. Later applications using similar designs used this approach (e.g., [16]). However, as we have shown here, there is a way to estimate those parameters using information from siblings-in-law of monozygotic and dizygotic twins. The key challenge (other than attaining sufficient statistical power) is that the variance of the sorting factor is unknown, which precludes the estimation of all relevant paths simultaneously. This is the same problem of identifying the scale that faces any latent variable model, and the solution is also the same: Either constrain the variance of the sorting factor to have unit variance (or some other constant) or fix one of the pathways – usually $\tilde{e}$ – and rescale afterwards.

#### Supplementary Note 2: The iAM-ACE model

The model in **Supplementary Figure 3** is identified (assuming e.g., $Var(S) = 1$) and it can be estimated with a structural equation model. In other words, twins-in-law can inform on the induced correlation in causes without having to assume direct assortment and while remaining agnostic to what associated trait(s) are undergoing assortment.

Most data sources that include twins and their partners will have data on both twins' partners. The full iAM-ACE model accommodates this by including two sets of partners linked together via a twinship or sibship (**Supplementary Figure 4**). This can increase statistical power by giving up to twice as many partners and in-law relations to analyse (for example, both the Partner 1 ↔ Twin 2 and Twin 1 ↔ Partner 2 dyads are twins-in-law). Additionally, including both partners of twins will yield a new type of relation into the model, namely that of the co-twins-in-law (The Partner 1 ↔ Partner 2 relation). This can further increase statistical power, or – as we will see below – be used to estimate other parameters. Finally, by adding twin-shared environmental factors (denoted $T$), one can readily include full siblings and their partners into the model without having to assume that the environments of twins and siblings are equally shared. Using full siblings in addition to twins can increase statistical power if available in sufficient numbers. The expected observed covariances between the observed variables are listed in **Supplementary Table 1**.

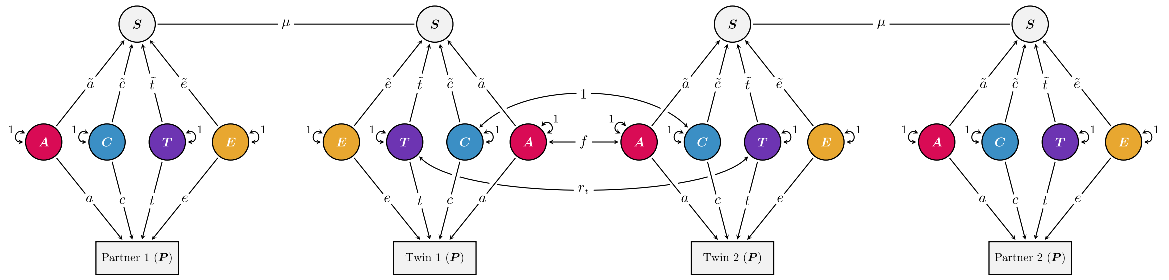

**Supplementary Figure 4** The iAM-ACE model. Differences in the observed, focal phenotype (denoted $P$) are thought to result from additive genetic factors ($A$), sibling-shared environmental factors ($C$), twin-shared environmental factors ($T$), and non-shared environmental factors ($E$). Their effects on the focal phenotype are denoted $a, c, t$, and $e$, respectively. The same factors also influence an unobserved sorting factor ($S$), with effects denoted $\tilde{a}, \tilde{c}, \tilde{t}$, and $\tilde{e}$, respectively. Partner similarity are thought to arise from assortative mating on the sorting factor, denoted with the co-path coefficient $\mu$. To be able to estimate the model, the variance of the sorting factor must be constrained, for example by fixing $e = \tilde{e}$). Some of the factors are correlated for twins (or siblings): Sibling-shared environmental factors are perfectly correlated across twin and sibling pairs, whereas the correlation in twin-shared environmental factors depend on relation (monozygotic and dizygotic twins: $r_t = 1$, ordinary full siblings: $r_t = 0$). Additive genetic factors are perfectly correlated across monozygotic twin pairs ($f = 1$), whereas for dizygotic twins and full siblings, the correlation is $f = (1 + \mu\tilde{a}^2)/2$ (assuming intergenerational equilibrium).

Twin-shared environments appear to be important for educational attainment[16]. The importance of twin-shared environments on the focal phenotype (denoted $t$) can be estimated by comparing correlations between dizygotic twins (where $r_t = 1$) and ordinary full siblings (where $r_t = 0$). They have the same genetic relatedness, meaning any increased similarity between dizygotic twins must be attributed to environmental factors shared by twins but not by siblings. By the same logic, the correlations between dizygotic twins-in-law and ordinary siblings-in-law provide the necessary information to estimate its importance for the sorting factor (denoted $\tilde{t}$).

**Supplementary Table 1**: Expected covariances in the iAM-ACE model (path specification)

| Description | Shorthand | Equation |
|---|---|---|
| Other terms: | | |
| Genotypic twin/sibling correlation<br>Twin 1 (A) ↔ Twin 2 (A) | $f$ | $f_{MZ} = 1;\ f_{DZ} = f_{FS} = \frac{1+\mu\tilde{a}^2}{2}$ |
| Twin-shared twin/sibling correlation<br>Twin 1 (T) ↔ Twin 2 (T) | $r_t$ | $r_{t_{MZ}} = r_{t_{DZ}} = 1;\ r_{t_{FS}} = 0$ |
| Covariance between focal phenotype and sorting factor<br>P ↔ S | $\sigma_{PS}$ | $a\tilde{a} + c\tilde{c} + t\tilde{t} + e\tilde{e}$ |
| Observed (co)variances: | | |
| Variance<br>Focal Phenotype, *P* | $\sigma_P^2$ | $a^2 + c^2 + t^2 + e^2$ |
| Partners<br>Partner 1 ↔ Twin 1; Partner 2 ↔ Twin 2 | $(\rho_{partners})$ | $\mu\sigma_{PS}^2$ |
| Twins / Siblings<br>Twin 1 ↔ Twin 2 | $(\rho_{twins})$ | $fa^2 + c^2 + r_t t^2$ |
| Twins-in-law<br>Partner 1 ↔ Twin 2; Twin 1 ↔ Partner 2 | $(\rho_{inlaw})$ | $\mu\sigma_{PS}(af\tilde{a} + c\tilde{c} + tr_t\tilde{t})$ |
| Co-twins-in-law<br>Partner 1 ↔ Partner 2 | | $\mu^2\sigma_{PS}^2(f\tilde{a}^2 + \tilde{c}^2 + r_t\tilde{t}^2)$ |

Model requires sorting factor to be constrained (e.g., by fixing $e = \tilde{e}$ or by constraining $Var(S) = \tilde{a}^2 + \tilde{c}^2 + \tilde{t}^2 + \tilde{e}^2 = 1$). If the focal phenotype is standardized ($\sigma_P^2 = 1$), the covariances can be interpreted as correlations.

#### Sub-models in the iAM-ACE model

The iAM-ACE model can be constrained to model specific types of assortative mating, which can be tested for significant differences in fit against the full model. For example, direct assortment can readily be tested by constraining $a = \tilde{a}$, $c = \tilde{c}$, $t = \tilde{t}$, and $e = \tilde{e}$, which in turn means that $\sigma_{PS} = \sigma_P^2 = \sigma_S^2$. This will give three degrees of freedom for the log likelihood test (assuming twin-shared environments are included).

Another sub-model is direct assortment with measurement error, where $\tilde{e}$ is freely estimated while all other paths to the sorting factor are constrained to equal the corresponding path leading to the focal phenotype ($\tilde{a} = a; \tilde{c} = c; \tilde{t} = t;\ \tilde{e} \neq e$). Note that this leads to different variances for the sorting factor and the focal phenotype, meaning that $a$ and $\tilde{a}$ may still be different after standardization (i.e., after rescaling to unit variance), but $\tilde{a}$, $\tilde{c}$, and $\tilde{t}$ will be inflated to the same degree.

It is also possible to model pure social homogamy ($\tilde{a} = 0; \tilde{c} = 1; \tilde{t} = 0, \tilde{e} = 0$) and pure genetic homogamy ($\tilde{a} = 1; \tilde{c} = 0; \tilde{t} = 0; \tilde{e} = 0$), although these are unlikely to be accurate descriptions of reality.

#### Alternative specification: Variance components versus path coefficients

The model can be parameterised primarily with path specification (where the model estimates path coefficients as in **Supplementary Figure 4**) or with variance specification (where the model estimates variance components as in **Supplementary Figure 5**). The two approaches are mathematically equivalent, but variance specification is often preferred due to superior error rate performance. When estimating variance components, the paths going to the focal phenotype are implicitly fixed to 1, whereas all but one of the paths going to the sorting factor can be freely estimated. The interpretation of $\tilde{a}$, $\tilde{c}$, $\tilde{t}$, and $\tilde{e}$ will here be slightly different. Rather than indicating the effect of the corresponding factor, it now indicates the relative importance of the corresponding factor on the sorting factor compared to the focal phenotype (i.e., weights). **Supplementary Table 2** lists the expected covariances for the iAM-ACE model using variance specification.

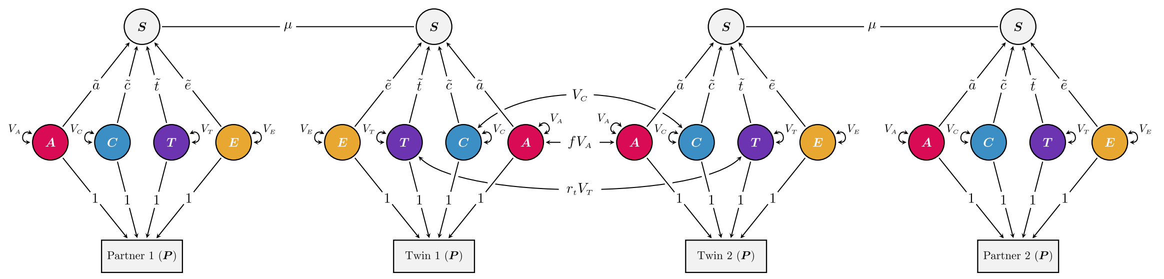

**Supplementary Figure 5** The iAM-ACE model with variance components instead of path coefficients

**Supplementary Table 2**: Expected covariances in the iAM-ACE model (variance specification)

| Description | Shorthand | Equation |
|---|---|---|
| <u>Other terms:</u> | | |
| Genotypic twin/sibling correlation<br>Twin 1 (A) ↔ Twin 2 (A) | $f$ | $f_{MZ} = 1;\ f_{DZ} = f_{FS} = \frac{1+\mu\tilde{a}^2 V_A}{2}$ |
| Twin-shared twin/sibling correlation<br>Twin 1 (T) ↔ Twin 2 (T) | $r_t$ | $r_{t_{MZ}} = r_{t_{DZ}} = 1;\ r_{t_{FS}} = 0$ |
| Covariance between focal phenotype and sorting factor<br>P ↔ S | $\sigma_{PS}$ | $V_A\tilde{a} + V_C\tilde{c} + V_T\tilde{t} + V_E\tilde{e}$ |
| <u>Observed (co)variances:</u> | | |
| Variance<br>Focal Phenotype, *P* | $\sigma_P^2$ | $V_A + V_C + V_T + V_E$ |
| Partners<br>Partner 1 ↔ Twin 1; Partner 2 ↔ Twin 2 | $(\rho_{partners})$ | $\mu\sigma_{PS}^2$ |
| Twins / Siblings<br>Twin 1 ↔ Twin 2 | $(\rho_{twins})$ | $fV_A + V_C + r_t V_T$ |
| Twins-in-law<br>Partner 1 ↔ Twin 2; Twin 1 ↔ Partner 2 | $(\rho_{inlaw})$ | $\mu\sigma_{PS}(fV_A\tilde{a} + V_C\tilde{c} + r_t V_T\tilde{t})$ |
| Co-twins-in-law<br>Partner 1 ↔ Partner 2 | | $\mu^2\sigma_{PS}^2(fV_A\tilde{a}^2 + V_C\tilde{c}^2 + r_t V_T\tilde{t}^2)$ |

Model requires sorting factor to be constrained (e.g., by fixing $\tilde{e} = 1$ or by constraining $Var(S) = V_A\tilde{a}^2 + V_C\tilde{c}^2 + V_T\tilde{t}^2 + V_E\tilde{e}^2 = 1$). If the focal phenotype is standardized ($\sigma_P^2 = 1$), the covariances can be interpreted as correlations.

#### Limitations and assumptions

The iAM-ACE model comes with many of the same assumptions as the classical twin model such as the equal environments assumption and no gene-environment interactions. However, unlike the classical twin model, the iAM-ACE model does not assume random mating, nor is it restricted to assuming direct assortment. However, to account for indirect assortative mating, two assumptions must be made.

First, the iAM-ACE model assumes that the observed partner correlation is fully attributable to assortative mating. If other causes of partner similarity are at play, such as convergence or social stratification, then the results may be inaccurate. The consequences of convergence may depend on the exact nature of the causal process, but in general, convergence will increase partner similarity with minimal effect on sibling and sibling-in-law similarity, thereby mimicking idiosyncratic homogamy (i.e., $\tilde{e}$). Social stratification and inbreeding will generally increase similarity among twins-in-law regardless of zygosity, meaning it will tend to mimic social homogamy (i.e., $\tilde{c}$). However, the other parameters may be slightly inaccurate because the model will, for example, overestimate the gene-environment correlations across partners. Social stratification may be estimated using the additional degree of freedoms afforded by co-siblings-in-laws (see description on page 15).

The second assumption about partner similarity concerns the history of assortment. If assortative mating has been occurring over many generations, the genotypic correlation between full siblings will have increased and stabilized at an equilibrium[8]. The analyses in this paper assume equilibrium. However, if the current generation was the first to assort, the genotypic correlation between siblings will be as under random mating. If assortative mating has been ongoing for several generations but not yet reached equilibrium, the genotypic correlation between siblings will be somewhere in between (although it will approach expectations under equilibrium after just a few generations, and be closer to equilibrium than random mating after only 1 generation[8]).

### Supplementary Note 3: Extensions of the iAM-ACE model

#### Sibling-shared mate preferences (i.e., cross-trait assortment)

Lindon Eaves has shown that if siblings or twins have correlated mate preferences, and these preferences are not perfectly correlated with their own trait value, then this may inflate the observed correlation between their respective partners[17]. This will be the case regardless of whether the preferences are active choices or merely imposed because of constraints in the environment. This will not affect correlations between partners or twins-in-law but will increase correlations between co-siblings-in-law (i.e., Partner 1 ↔ Partner 2). The increased correlations may bias the results unless accounted for.

Here, mate preferences can be conceived as a separate trait, making this a form of cross-trait assortative mating. Incorporating multivariate assortment across multiple observed phenotypes is beyond the scope of this paper. Here, we simply show how it affects familial correlations for a single observed trait and show how to account for it in the iAM-ACE model. **Supplementary Figure 6** includes a secondary sorting factor (denoted $S'$), which may indicate individual differences in mate preferences orthogonal to (or conditional on) a person's own trait value. We suppose that partners are matching on both the primary and secondary sorting factors, indicated with four co-paths for each mating process. Because the same mating process is now indicated with several co-paths, we must add the path tracing rule that a chain cannot include multiple co-path coefficients stemming from the same mating process (i.e., linking the same pair of partners). Co-paths and variables irrelevant for the observed correlations are greyed out.

To the extent that such cross-trait assortment affects the mating process, the preferences of one person will be associated with the trait of their partner (indicated with the co-path denoted $\mu'$). To the extent that twins or siblings have similar preferences (denoted $r_{S'}$), they will tend to end up with partners having similar trait values over and above the effects of assortment on the primary sorting factor. The co-siblings-in-law correlation will therefore increase by $\sigma_{PS}^2 \mu'^2 r_{S'}$. It is not possible to distinguish $\mu'^2$ and $r_{S'}$ when looking at a single trait, but it is possible to estimate a model where their product (called, say, $\zeta$) is estimated separately across sibship types, thus removing any potential bias this may have on the remaining parameters.

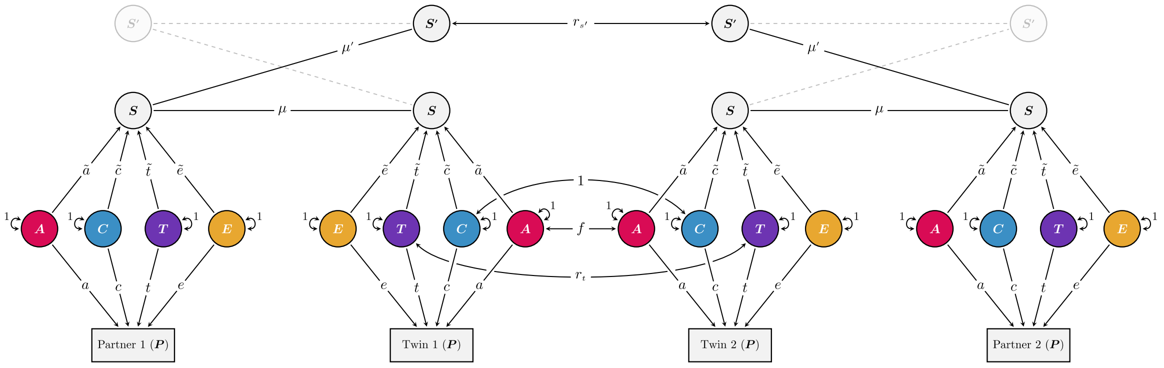

**Supplementary Figure 6** The iAM-ACE model with a secondary sorting factor (denoted $S'$) orthogonal to the primary sorting factor. The secondary sorting factor includes mate preferences or any other trait undergoing cross-trait assortment with the primary sorting factor. This is indicated with a co-path (denoted $\mu'$) going from the primary sorting factor to their partner's secondary sorting factor (irrelevant paths are greyed out). Such cross-trait assortment may inflate the correlation between co-siblings-in-law (Partner 1 ↔ Partner 2) to the extent that the secondary sorting factor is correlated for siblings or twins ($r_{S'}$).

#### Social Stratification

If partner similarity arises because of social stratification rather than assortative mating, then this should presumably affect similarity between all family members in **Supplementary Figure 4** equally. The added degrees of freedom offered by including co-siblings-in-law makes it possible to estimate the effects of social stratification (provided sibling-shared mate preferences are not included). In **Supplementary Figure 7**, social stratification (denoted $Q$) is included as an additional environmental factor that influence the observed focal phenotypes equally for all family members but are completely unrelated to the sorting factor.

Here, all observed correlations are increased with $q^2$, which reduces the relative difference between the siblings-in-law correlation (e.g., Partner 1 ↔ Twin 2) and co-siblings-in-law correlation (Partner 1 ↔ Partner 2). As this difference is likely to be very small, the power to distinguish social stratification and assortative mating is likely poor.

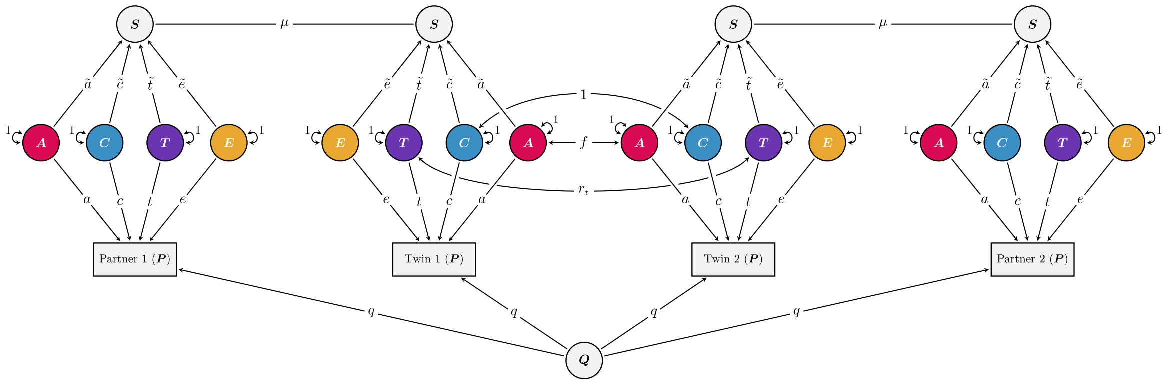

**Supplementary Figure 7** The iAM-ACE model with social stratification

#### Other extensions

Numerous other extensions are imaginable. One weakness with the iAM-ACE model is that it cannot estimate gene-environment correlations. In classical twin models, passive gene-environment correlations will simply look like sibling-shared environmental effects and is as such not a very important limitation. However, gene-environment correlations will entangle the genetic and social consequences of assortative mating[8], meaning, for example, that the genetic consequences may be underestimated unless it is included in the model. To get around this, it is possible to include it in the iAM-ACE model as a fixed value (i.e., assume it is, say .20 rather than .00). An alternative is to include children of twins and siblings and use them to investigate causes of intergenerational transmission (which will yield an estimate of the gene-environment correlation). This will also take care of a second weakness with the iAM-ACE model, namely that it is not as informative on the intergenerational consequences of indirect assortative mating. In the next section, we describe a children-of-twins-and-siblings model that can account for indirect assortative mating.

#### Supplementary Note 4: The iAM-COTS model

In **Supplementary Figure 8**, the iAM-ACE model is extended with observations on two children per partnership. The model is now a variant of the children-of-twins-and-siblings model (aka., COTS model), extended with partners and multiple children[18-20].

To differentiate factors that operate on the different generations, each variable and its effect have been subscripted with 1 if they operate on the parental phenotype (i.e., if they were included in the original iAM-ACE model) and 2 if they operate exclusively on the offspring phenotype. That is, the model does not strictly assume that the parental and offspring phenotypes are the same. The offspring phenotype is decomposed similarly to the parental phenotype, albeit with no twin-shared environments (i.e., with additive genetic, sibling-shared environmental, and non-shared environmental factors). The additive genetic factor is split into the component that is associated with the parental phenotype ($A_1'$) and the component that is unique to the offspring phenotype ($A_2$). The offspring genetic factor associated with the parental phenotype is a function of the parental genetic factors and recombination variance (denoted $k$, equal to $1 - f$ in intergenerational equilibrium). The other genetic factor, $A_2$, is correlated between siblings (.50) and cousins ($q$). The genotypic correlation between cousins will depend on whether their parents are monozygotic twins or not ($q_{MZ} = .25; q_{DZ} = q_{FS} = .125$).

The sibling-shared environmental influences are also split into that which is associated with the parental phenotype via some form of environmental transmission ($F$) and that which is unique to the offspring generation ($C_2$). The model includes two forms of environmental transmission: direct phenotypic transmission where the parental phenotype influences the offspring phenotype directly ($p$), and passive environmental transmission where the sibling-shared environmental factor that influenced the parents also influence the offspring ($c_1'$). If both genetic and environmental transmission are non-zero, the effects of $A_1'$ and $F$ can become correlated (i.e., a gene-environment correlation). The model can use this gene-environment correlation as a best guess for what the gene-environment correlation is in the parent-generation, such that the correlation between $C_1$ and $A_1$, denoted $\omega$, is constrained to equal the correlation between $(A_1' + A_2)$ and $(F + C_2)$.

Different forms of environmental transmission have been discussed elsewhere[21], and outstanding problems are beyond the scope of this paper. For example, phenotypic transmission can be indirect via another trait or set of traits, similar to how assortment can be indirect[6]. This may lead to different expectations than those explored here. Similarly, passive environmental transmission, where the parent-offspring correlation is confounded by shared environmental influences, can also be specified in multiple different ways, with slightly different expectations. For example, expectations depend on whether environmental factors are separated into shared and non-shared components (where passive environmental transmission are imagined as $C \rightarrow C$), or simply treated as one factor with a non-unit correlation between family members (where passive environmental transmission are imagined as $E \rightarrow E$, such as in [21] and in the supplement of [8]). The key difference concerns which sources of new environmental variance are transmitted to the next generation, which are necessary for transmitted variance to be stable across multiple generations[1]. We do not have enough information to distinguish intergenerational persistence of environmental factors, generation-specific sources of variance on transmitted factors, and the effect of those factors on the offspring phenotype. These are important avenues to explore, theoretically and empirically, but it falls beyond the scope of this paper. Here, we have opted for a variant previously used in children-of-twins models where the parental shared environmental factors ($C_1$) directly influence the offspring phenotype ($C \rightarrow P$)[18,19]. (In the path diagram, these paths have been gathered into a latent variable $F$ to reduce clutter, together with the paths from the parental phenotypes, although the expectations are equivalent to paths going directly to the offspring phenotype).

The expected covariances along with some useful shorthand are presented in **Supplementary Table 3**. The key correlation of interest is that between parents and offspring:

$$Cov(Parent, Offspring) = \frac{(a_1 + c_1\omega)a'_1}{2} + (c_1 + a_1\omega)c'_1 + \sigma_p^2 p + \sigma_{PS}\mu\lambda_{SO} \quad (9)$$

Here, the first component (**red**) represents that which is attributable to genetic transmission, the second component (**blue**) represents that which is attributable to passive environmental transmission, the third component (**yellow**) represents that which is attributable to direct phenotypic transmission, and finally the fourth component (**green**) represents the inflation due to effects via the other parent. ($\lambda_{SO}$ represents the covariance between the parental sorting factor and the offspring phenotype). It is possible to solve for these different components by comparing the parent-offspring covariance with that of the avuncular covariance across zygosity:

$$Cov(Uncle\ or\ Aunt, Niece\ or\ Nephew) = \frac{(fa_1 + c_1\omega)a'_1}{2} + (c_1 + a_1\omega)c'_1 + \delta_{PP}p + \delta_{PS}\mu\lambda_{SO} \quad (10)$$

Here, $\delta$ refers to the covariance between siblings or twins ($\delta_{PP}$ for the phenotypic covariance, and $\delta_{PS}$ for the covariance between one twin's phenotype and the other twin's sorting factor). If we momentarily disregard assortative mating (i.e., assume $\mu = 0$), then we can easily see how these two covariances are informative on the nature of the parent-offspring covariance. For monozygotic twin families, where $f = 1$, offspring are as related to their parents as they are to their parent's twin. Here, the first two components in **Equation 9 and 10** are equal, and the only thing that differs is the consequence of direct phenotypic transmission (**yellow**). Consequently, if the parent-offspring correlation is larger than the avuncular correlation in monozygotic twin families, then this would imply phenotypic transmission operating within the nuclear family (assortative mating notwithstanding). If, on the other hand, the parent-offspring correlation would equal the avuncular correlation, then this would imply passive forms of transmission – either genetic or environmental – that are shared across the extended family. The effects of assortative mating are a bit less straightforward to glean

[1] If transmitted environmental factors ($F_o$) are fully determined by paternal ($F_p$) and maternal ($F_m$) transmitted environmental factors, then the variance will not be stable across multiple generations unless the paths from parent environments to offspring environment are exactly $\sqrt{0.5}$. However, variance can be intergenerationally stable with intergenerational paths less than $\sqrt{0.5}$ if new sources of variance are also transmitted to the next generation (similar to how recombination variance enables intergenerationally stable genetic variance despite the intergenerational paths being 0.5 (which is less than $\sqrt{0.5}$).

intuitively but can be accounted for when estimating the model. By comparing the avuncular correlation across zygosity (i.e., across various levels of $f$), it is possible to disentangle passive genetic transmission (**red**) from passive environmental transmission (**blue**). In the extreme cases, pure genetic transmission would result in the avuncular correlation in monozygotic twin families to be about twice as large as in dizygotic twin and full sibling families (assuming no assortative mating), whereas pure passive environmental transmission would result in the same avuncular correlation across zygosity.

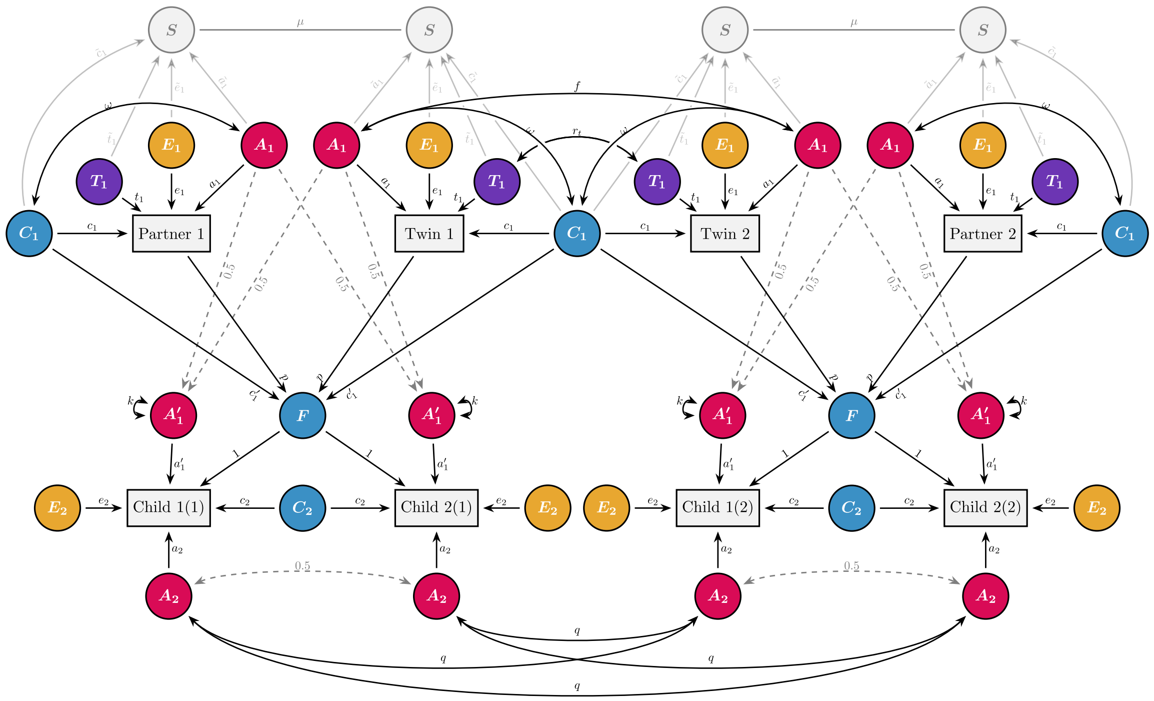

**Supplementary Figure 8** The iAM-COTS model. The model expands the iAM-ACE model to include their children, which allows decomposing the parent-offspring correlation into genetic and environmental pathways, which, in turn, estimates gene-environment correlations.

**Supplementary Table 3: Equations implied by the iAM-COTS model in Supplementary Figure S8**

| Name | Shorthand | Equation | Note |
|---|---|---|---|
| **Useful Shorthands** | | | |
| ***Variance Components in Parent generation*** | | | |
| Additive genetic variance | $V_{A1} =$ | $a_1^2$ | |
| Sibling-shared environmental variance | $V_{C1} =$ | $c_1^2$ | |
| Twin-shared environmental variance | $V_{T1} =$ | $t_1^2$ | $r_{t_{MZ}} = r_{t_{DZ}} = 1; r_{t_{FS}} = 0$ |
| Gene-Environment covariance | $V_{rGE1} =$ | $2\omega a_1 c_1$ | |
| Non-shared environmental variance | $V_{E1} =$ | $e_1^2$ | |
| ***Variance Components in Offspring generation*** | | | |
| Additive genetic variance associated with parental phenotype | $V_{A1P} =$ | $a'^2_1(f_{FS} + k)$ | $f_{FS} + k = 1$ |
| Family variance associated with parental phenotype | $V_F =$ | $2\sigma_p^2 p^2 + 2c'^2_1 + 4p(c_1 + a_1\omega)c'_1 + 2\mu(c'_1(c_1 + a_1\omega) + \sigma_{PS}p)^2$ | *Both direct and passive environmental effects* |
| Additive genetic variance independent of parental phenotype | $V_{A2} =$ | $a_2^2$ | |
| Shared environmental variance independent of parental phenotype | $V_{C2} =$ | $c_2^2$ | |
| Gene-Environment covariance | $V_{rGE2} =$ | $2p(a_1 + c_1\omega)a'_1 + 2a'_1\omega c'_1 + 2a'_1(\tilde{a}_1 + \tilde{c}_1\omega)\mu((\tilde{c}_1 + \tilde{a}\omega)c'_1 + \sigma_{PS}p)$ | |
| Unique environmental variance | $V_{E2} =$ | $e_2^2$ | |

**Supplementary Table 3: Equations implied by the iAM-COTS model in Supplementary Figure S8**

| Name | Shorthand | Equation | Note |
|---|---|---|---|
| ***Other useful Shorthands*** | | | |
| Correlation between P and S | $\sigma_{PS} =$ | $(\tilde{a}_1 + \tilde{c}_1\omega)a_1 + (\tilde{c}_1 + \tilde{a}_1\omega)c_1 + t_1\tilde{t}_1 + e_1\tilde{e}_1$ | *Focal phenotype - Sorting factor correlation* |
| Sibling Correlation between P and S | $\delta_{PS} =$ | $(f\tilde{a}_1 + \tilde{c}_1\omega)a_1 + (\tilde{c}_1 + \tilde{a}_1\omega)c_1 + \tilde{t}_1 r_t t_1$ | *Correlation between focal phenotype and their siblings sorting factor* |
| Sibling Correlation between S and S | $\delta_{SS} =$ | $f\tilde{a}_1^2 + \tilde{c}_1^2 + 2\tilde{a}_1\omega\tilde{c}_1 + r_t\tilde{t}_1^2$ | *Correlation between siblings' sorting factors* |
| Genotypic correlation between full siblings | $f_{FS} =$ | $\frac{1 + \mu(\tilde{a}_1 + \tilde{c}_1\omega)^2}{2}$ | $f_{FS} = f_{DZ}$ |
| Focal Phenotype - Offspring Covariance w/o assortative mating | $\lambda_{PO} =$ | $\frac{(a_1 + c_1\omega)a'_1}{2} + (c_1 + a_1\omega)c'_1 + \sigma_p^2 p$ | *Correlation between parental (focal) phenotype and offspring phenotype* |
| Sorting Factor - Offspring Covariance w/o assortative mating | $\lambda_{SO} =$ | $\frac{(\tilde{a}_1 + \tilde{c}_1\omega)a'_1}{2} + (\tilde{c}_1 + \tilde{a}_1\omega)c'_1 + \sigma_{PS} p$ | *Correlation between parental sorting factor and offspring phenotype* |
| **Cells in Covariance Matrix** | | | |
| ***Variances*** | | | |
| Observed Parental Phenotype | $\sigma_p^2 =$ | $V_{A1} + V_{C1} + V_{T1} + V_{rGE1} + V_{E1}$ | |
| Offspring Phenotype | $\sigma_o^2 =$ | $V_{A1P} + V_F + V_{A2} + V_{C2} + V_{rGE2} + V_{E2}$ | |
| ***Covariances in Parent Generation*** | | | |
| Siblings/Twins | $\delta_{PP} =$ | $fV_{A1} + r_t V_{T1} + V_{rGE1} + V_{C1}$ | |
| Partners | | $\sigma_{PS}^2\mu$ | |

**Supplementary Table 3: Equations implied by the iAM-COTS model in Supplementary Figure S8**

| Name | Shorthand | Equation | Note |
|---|---|---|---|
| Siblings-in-law | | $\sigma_{PS}\mu\delta_{PS}$ | |
| Co-siblings-in-law | | $\sigma_{PS}^2\mu^2\delta_{SS}$ | |
| ***Intergenerational Covariances*** | | | |
| Parent-Offspring | | $\lambda_{PO} + \sigma_{PS}\mu\lambda_{SO}$ | |
| Avuncular | | $\frac{(fa_1 + c_1\omega)a'_1}{2} + (c_1 + a_1\omega)c'_1 + \delta_{PP}p + \delta_{PS}\mu\lambda_{SO}$ | *The genetically related uncle/aunt* |
| Avuncular-in-law | | $\sigma_{PS}\mu\left(\left(\frac{(f\tilde{a}_1 + \tilde{c}_1\omega)a'_1}{2} + (\tilde{c}_1 + \tilde{a}_1\omega)c'_1 + \delta_{PS}p\right) + \delta_{SS}\mu\lambda_{SO}\right)$ | *The non-genetically related uncle/aunt* |
| ***Covariances in Offspring Generation*** | | | |
| Siblings | | $f_{FS}a'^2_1 + \frac{V_{A2}}{2} + V_F + V_{C2} + V_{rGE2}$ | |
| Cousins | | $\frac{fa'^2_1}{4} + qV_{A2} + \delta_{PP}p^2 + c'^2_1 + 2p(c_1 + a_1\omega)c'_1 + p(fa_1 + c_1\omega)a'_1 + a'_1\omega c'_1$ $+ 2\lambda_{SO}\mu\left(\frac{(f\tilde{a}_1 + \tilde{c}_1\omega)a'_1}{2} + (\tilde{c}_1 + \tilde{a}_1\omega)c'_1 + \delta_{PS}p\right)$ $+ \lambda_{SO}^2\mu^2\delta_{SS}$ | |

#### Supplementary Note 5: Simulations

In this section, we provide simulations that show that the iAM-ACE and iAM-COTS models can retrieve the simulated parameters based on observed phenotypes. For both models, we simulated the implied causal model, meaning we randomly simulated all exogenous sources of (co)variation before simulating the effects of that variation. The scripts and result files for the simulations are available at https://osf.io/dznbk/.

##### Simulating the iAM-ACE model

To test the iAM-ACE model, we simulated a set of twins (or siblings) and their partners. To accomplish this, we first simulated $A$, $C$, $T$, and $E$ as normally distributed random variables for each individual, with the desired correlations between twins/siblings (using `mvrnom` from the MASS package in R). The focal phenotypes and sorting factors were then calculated as a weighted sum of $A$, $C$, $T$, and $E$, using the desired path coefficients as weights. Prospective partners were handled in separate data frames, which were sorted on a combination of the sorting factor and random error before being joined together (where individuals on the same row were considered partners). The amount of random error allowed us to achieve the desired correlation between partners on the sorting factor.

We estimated the model on 1000 simulated samples of 3,000 monozygotic twin families, 3,000 dizygotic twin families, and 150,000 full sibling families. The true variance decomposition of the focal phenotype was constant across samples at $V_A$ = 50%, $V_C$ = 10%, $V_T$ = 5% and $V_E$ = 35%. For the sorting factor, we randomly and independently varied the true amount of genetic and social homogamy, from 0% to 50% variance explained in the sorting factor each (with the remaining variance being explained by non-shared environmental influences). We set the desired partner correlation on the focal phenotype to .50 and let the implied correlation on the sorting factor vary depending on the degree of genetic and social homogamy. If the random combination of parameters were impossible (e.g., because of an implied partner correlation above 1), a new set of random parameters were drawn. If the first attempt at model fitting did not converge satisfactorily in a given sample, we attempted to refit the model using `mxTryHard`. Of the 1000 models, 954 eventually converged satisfactorily. Samples where the model did not converge are not included in the remaining figures. One sample yielded very unsensible estimates ($V_A = 105\%, V_C = -44\%$), which we removed from the below figures.

**Supplementary Figure 9a** shows the results for the variance decomposition of the focal phenotype, which was simulated to be constant despite varying degrees of genetic and social homogamy. Each dot (within each parameter) represents one simulated sample. The large, black dots on the right of each parameter are the true, simulated parameter while the lines on the left cover 50% (thick portion) and 95% (thin portion) of the sample of estimated parameters. The grey, dashed line within each parameter is the median estimate across all samples. Sampling variation notwithstanding, the model is able to retrieve the simulated parameters adequately. The model is also able to estimate the partner correlation on the sorting factor (panel **b**) across varying simulated values. Panel **c** and **d** shows that the model can retrieve the simulated values of genetic and social homogamy, respectively. However, sampling variation is more pronounced for these values.

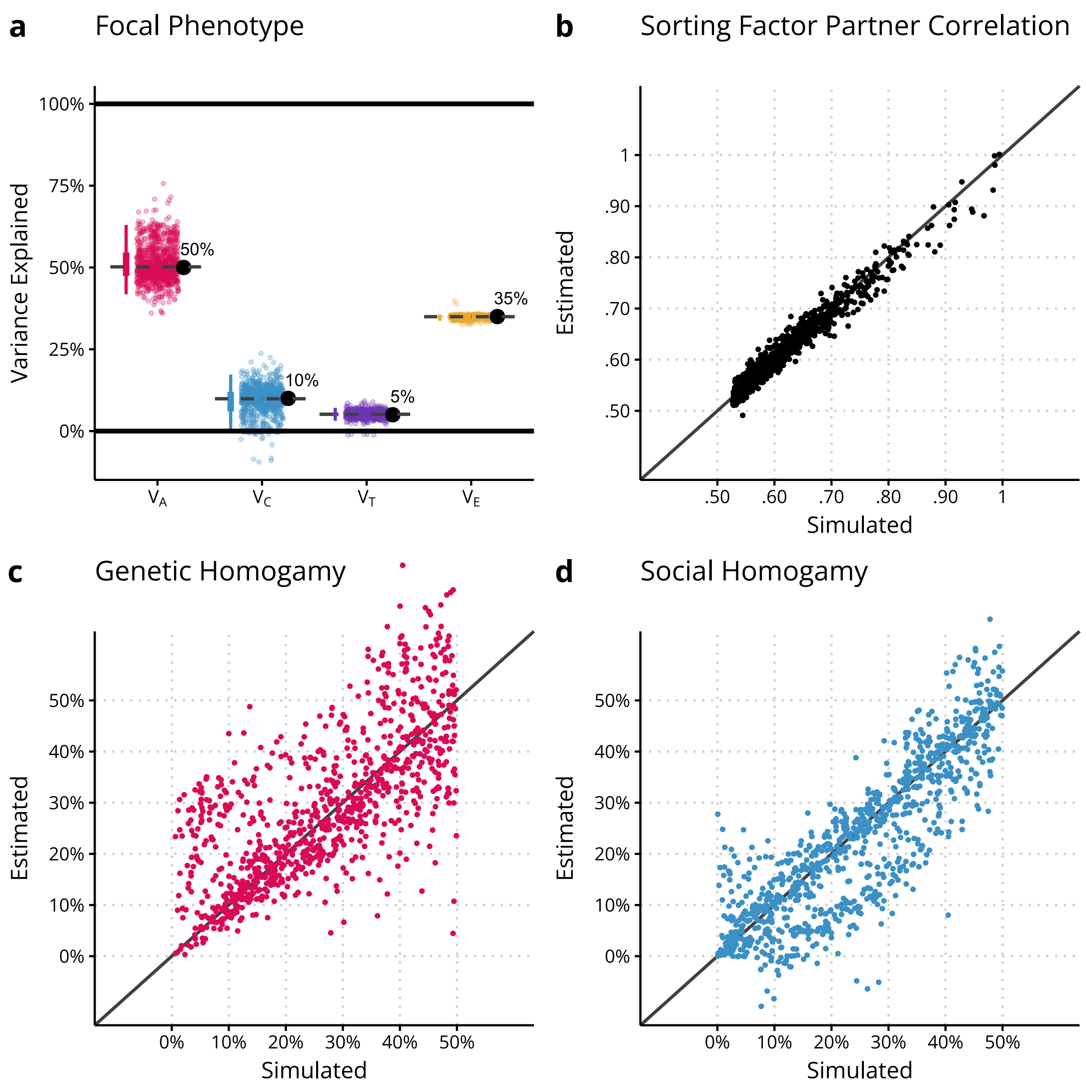

**Supplementary Figure 9** Results from estimating the iAM-ACE model on 953 simulated samples of 3,000 monozygotic twin families, 3,000 dizygotic twin families, and 150,000 full sibling families. (**a**) Variance decomposition of the focal phenotype. Each dot (within a given parameter) represents one simulated sample. The large, black dots on the right (or top) of each parameter are the true, simulated parameter while the lines on the left (or bottom) cover 50% (thick portion) and 95% (thin portion) of the sample of estimated parameters. The grey, dashed lines are the median estimates across all samples. (**b**) Scatterplot of the simulated partner correlation for the sorting factor against the model estimated partner correlation. (**c**) Scatterplot of the simulated amount of genetic homogamy (stated as variance explained in the sorting factor) against the estimated amount of genetic homogamy. (**c**) Scatterplot of the simulated amount of social homogamy (stated as variance explained in the sorting factor) against the estimated amount of social homogamy.

#### The iAM-COTS model

To test the iAM-COTS model, we expanded the above simulation by also simulating offspring. We again simulated exogeneous variables ($A$, $C$, $T$, and $E$) as normally distributed random variables (with correlations where appropriate, such as between $A$ and $C$ and between twins/siblings), and calculated the endogenous variables as described above. Offspring genotypes were simulated as a function of parental genotypes and random recombination variance. We did this for both the genetic factor associated with the parental phenotype ($A_1'$) and the genetic factor unique to the offspring phenotype ($A_2$). The offspring phenotypes were then calculated as implied in **Supplementary Figure 8**, namely as a weighted sum of the parental phenotype, the parental shared environments ($C_1$), the offspring genetic factors ($A'_1$ and $A_2$), and offspring-specific environmental factors ($C_2$ and $E_2$).

We again estimated the model on 1000 simulated samples of 3,000 monozygotic twin families, 3,000 dizygotic twin families, and 150,000 full sibling families. For these simulations, all true parameters were constant across samples (see large, black dots in **Supplementary Figure 10**). The observed partner correlation was set at .45, whereas the partner correlation on the sorting factor was simulated at .53. If the first attempt at model fitting did not converge satisfactorily in a given sample, we attempted to refit the model using `mxTryHard`. Of the 1000 models, all eventually converged satisfactorily.

**Supplementary Figure 10** and shows the results for the focal phenotype (panel **a**), the sorting factor (**b**), the offspring phenotype (**c**), and the parent-offspring correlation (**d**). As with the results for the iAM-ACE model, the iAM-COTS model is able to retrieve the simulated parameters adequately. The accuracy is, as expected, higher for the focal phenotype than the sorting factor. The sampling variation is also not symmetric for genetic and social homogamy (even though the median estimate is unbiased).

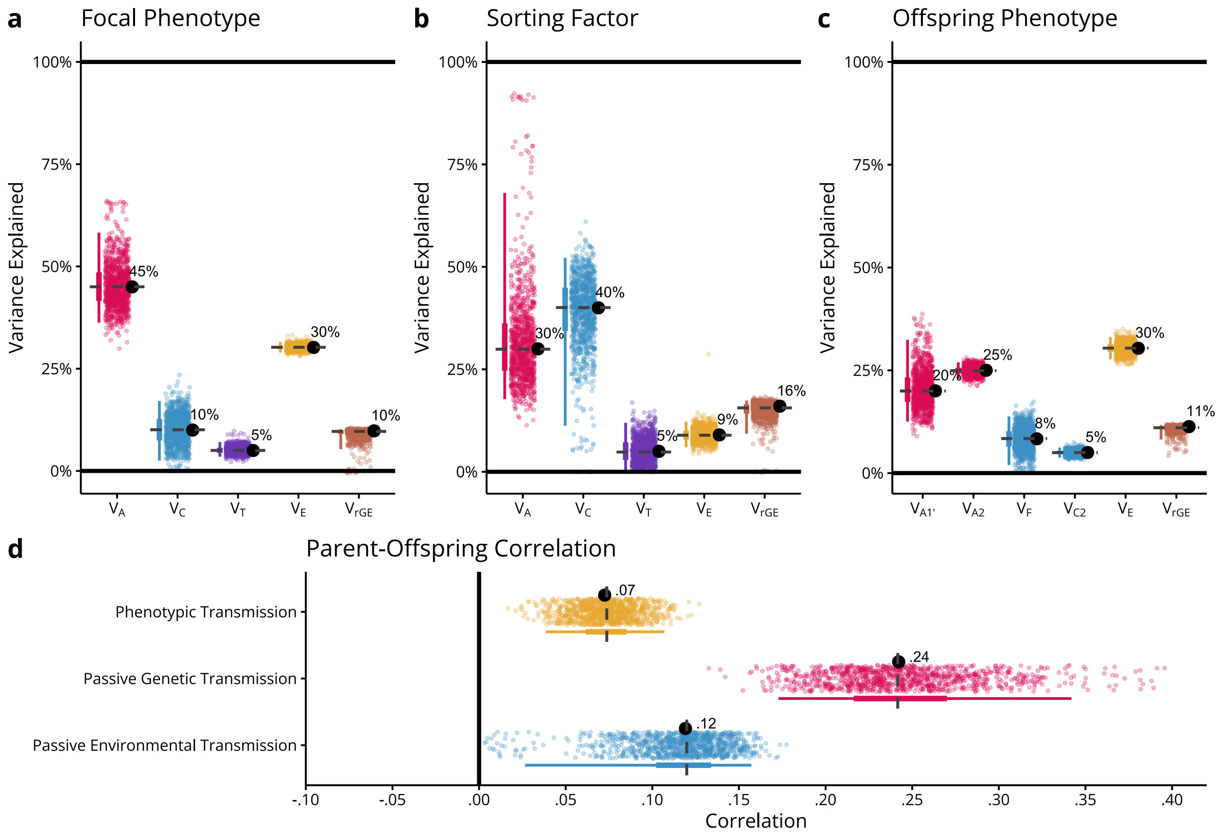

**Supplementary Figure 10** Results from estimating the iAM-COTS model on 1000 simulated samples 3,000 monozygotic twin families, 3,000 dizygotic twin families, and 150,000 full sibling families. Each dot (within a given parameter) represents one simulated sample. The large, black dots on the right (or top) of each parameter are the true, simulated parameter while the lines on the left (or bottom) cover 50% (thick portion) and 95% (thin portion) of the sample of estimated parameters. The grey, dashed lines are the median estimates across all samples.

### Supplementary Tables and Figures

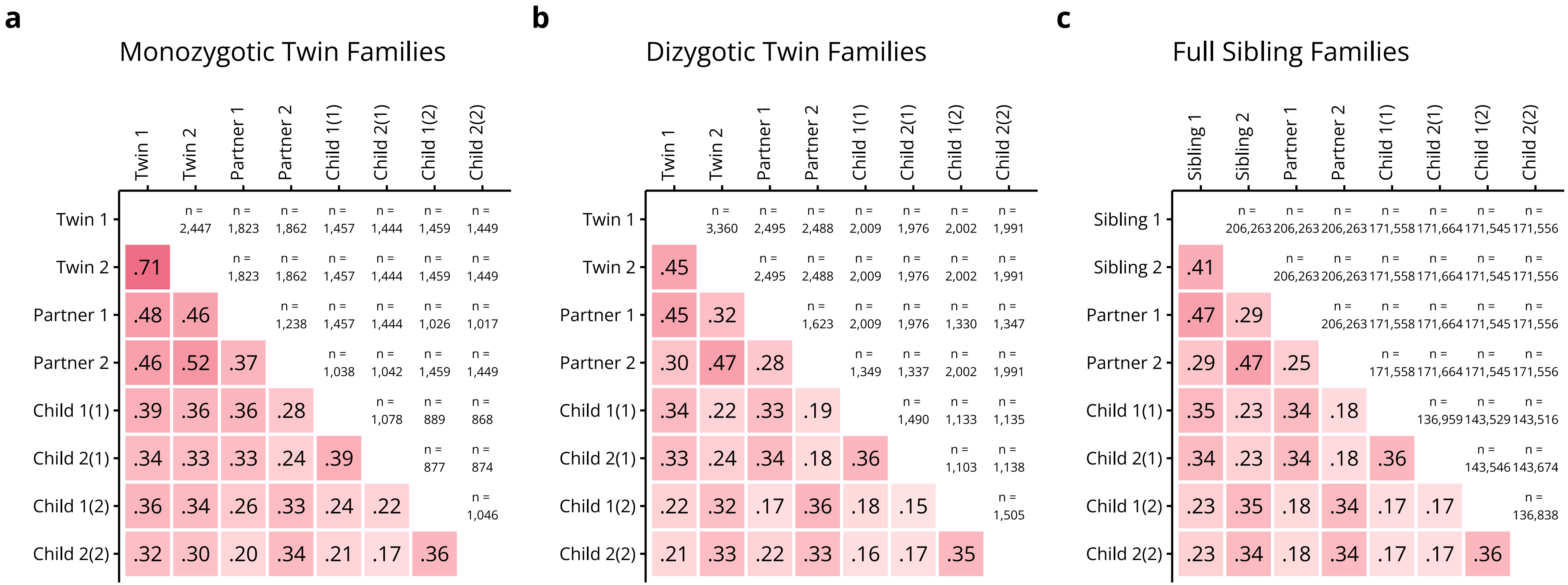

**Supplementary Figure 11** Correlation matrices and sample sizes for extended families of (**a**) monozygotic twins, (**b**) dizygotic twins, and (**c**) full siblings. Correlations are also available in a data file at https://osf.io/dznbk/.

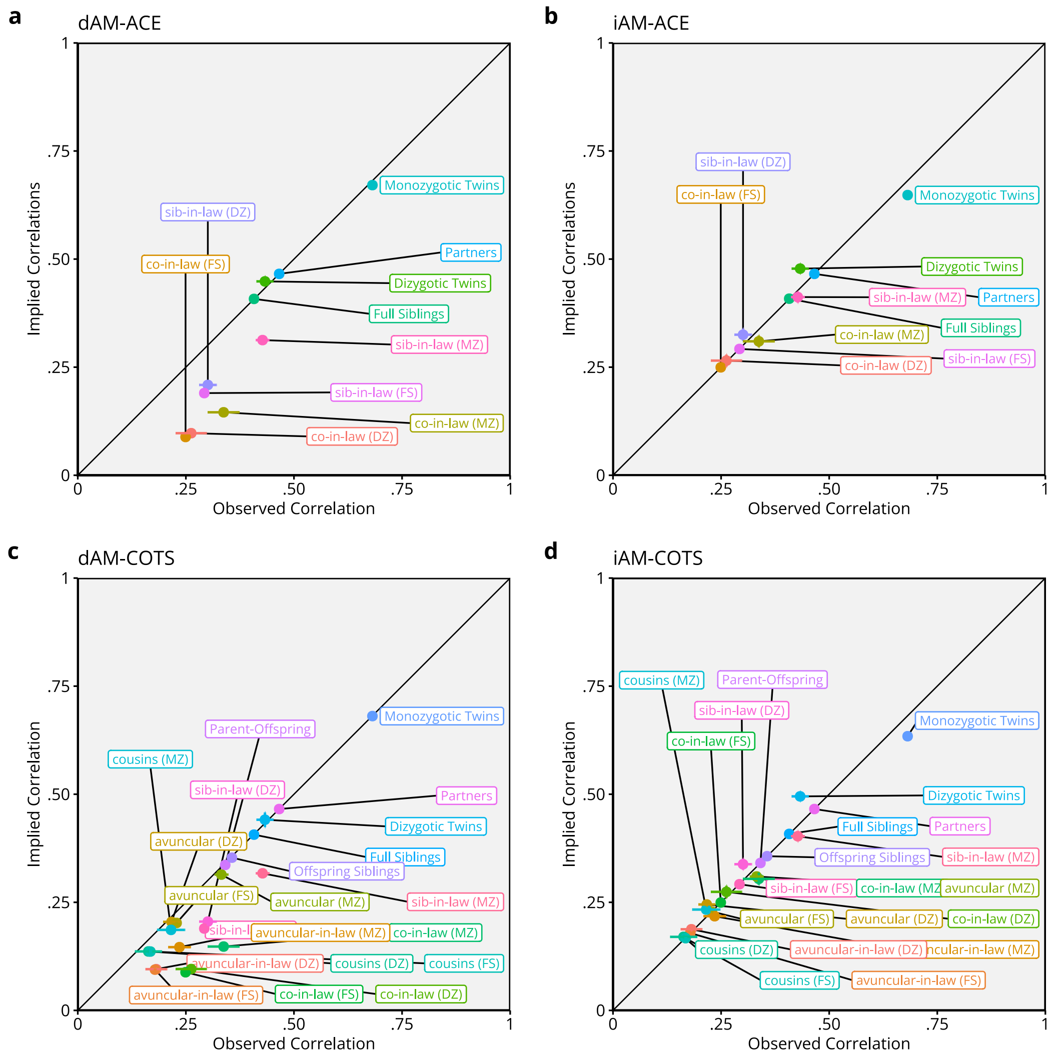

**Supplementary Figure 12** Comparisons of observed versus model-implied phenotypic correlations between family members in 212,070 extended families for models constrained to direct assortment (dAM, panel **a** and **c**) and for full models (iAM, panel **b** and **d**). Observed correlations are estimated using OpenMx where equivalent correlations (e.g., Partners in the different zygosity groups) are constrained to be equal. Implied correlations are extracted from the fitted models. Error bars are 95% Wald-type confidence intervals. Correlations are also available in data files at https://osf.io/dznbk/.

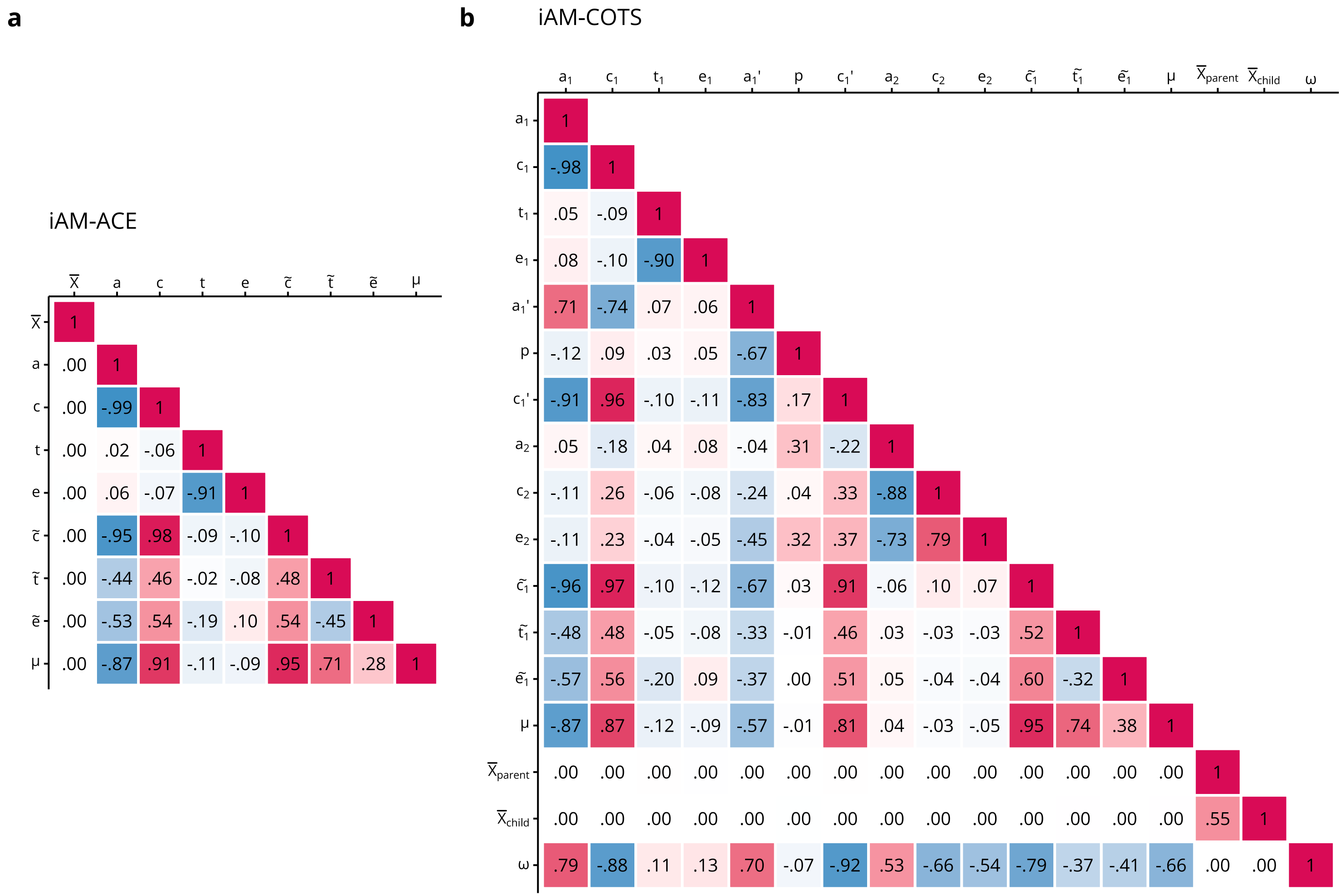

**Supplementary Figure 13** Correlations between maximum likelihood parameter estimates in the iAM-ACE (**a**) and iAM-COTS (**b**) models applied to 212,070 extended families. Colors are proportional to correlation (-1 = blue, +1 = red). There is no $\tilde{a}$ because variance of the sorting factor was fixed by fixing $\tilde{a} = a$. Alternative parameterizations may yield different correlations. The unstandardized variance-covariance matrices are available in a data file at https://osf.io/dznbk/.

**Supplementary Table 4: Correlations (95% Confidence Intervals)**

| Relation | Monozygotic | Dizygotic | Full Sibling |
|---|---|---|---|
| Siblings/Twins | .706<br>*(.696, .715)* | .450<br>*(.428, .470)* | .407<br>*(.404, .410)* |
| Partners | .504<br>*(.485, .523)* | .460<br>*(.442, .477)* | .465<br>*(.463, .467)* |
| Siblings-in-law | .462<br>*(.441, .483)* | .310<br>*(.286, .334)* | .292<br>*(.289, .295)* |
| Co-Siblings-in-law | .371<br>*(.332, .410)* | .274<br>*(.234, .312)* | .249<br>*(.245, .253)* |
| Parent-Offspring | .341<br>*(.321, .361)* | .333<br>*(.317, .350)* | .341<br>*(.339, .343)* |
| Avuncular | .340<br>*(.318, .362)* | .223<br>*(.200, .245)* | .228<br>*(.226, .231)* |
| Avuncular-in-law | .247<br>*(.216, .277)* | .183<br>*(.156, .211)* | .178<br>*(.176, .181)* |
| Siblings (Offspring) | .370<br>*(.339, .400)* | .355<br>*(.328, .381)* | .356<br>*(.353, .359)* |
| Cousins (Offspring) | .216<br>*(.179, .252)* | .161<br>*(.128, .194)* | .168<br>*(.165, .171)* |

**Supplementary Table 5: Testing parameters in the iAM-ACE model**

| Base | Comparison | Parameters | -2LL | df | ΔLL | Δdf | *p* |
|---|---|---|---|---|---|---|---|
| + Stratification | | 10 | 2,247,740 | 845,324 | | | |
| **+ Stratification** | **Full iAM-ACE model**[a] | **9** | **2,247,740** | **845,325** | **0.0** | **1** | $\mathbf{9.997 \times 10^{-1}}$ |
| Full iAM-ACE model | Direct Assortment + Measurement Error | 7 | 2,248,277 | 845,327 | 536.9 | 2 | $2.611 \times 10^{-117}$ |
| Full iAM-ACE model | Direct Assortment | 6 | 2,256,338 | 845,328 | 8,598.2 | 3 | 0 |
| Full iAM-ACE model | No Assortment ($\mu = 0$) | 5 | 2,359,308 | 845,329 | 111,567.6 | 4 | 0 |

[a]Reported parameters are from this model

**Note.** Model comparisons were evaluated using likelihood-ratio tests of nested models, calculated as differences in $-2LL$. Test statistics follow a one-sided $\chi^2$ distribution with $\Delta df$ degrees of freedom. No corrections were made for multiple comparisons

**Supplementary Table 6: Testing parameters in the iAM-COTS model**

| Base | Comparison | Parameters | -2LL | df | *Δ*LL | *Δ*df | *p* |
|---|---|---|---|---|---|---|---|
| **Full iAM-COTS model[a]** | | **17** | **4,090,739** | **1,545,428** | | | |
| Full iAM-COTS model | $a_1' = 0$ | 16 | 4,090,936 | 1,545,429 | 196.8 | 1 | $1.052 \times 10^{-44}$ |
| Full iAM-COTS model | $c_1' = 0$ | 16 | 4,090,752 | 1,545,429 | 12.8 | 1 | $3.492 \times 10^{-4}$ |
| Full iAM-COTS model | $p = 0$ | 16 | 4,090,786 | 1,545,429 | 46.4 | 1 | $9.651 \times 10^{-12}$ |
| Full iAM-COTS model | Direct Assortment + Measurement Error | 15 | 4,091,254 | 1,545,430 | 514.3 | 2 | $2.068 \times 10^{-112}$ |
| Full iAM-COTS model | Direct Assortment | 14 | 4,100,038 | 1,545,431 | 9,299.0 | 3 | 0 |

[a]Reported parameters are from this model

**Note.** Model comparisons were evaluated using likelihood-ratio tests of nested models, calculated as differences in $-2LL$. Test statistics follow a one-sided $\chi^2$ distribution with $\Delta df$ degrees of freedom. No corrections were made for multiple comparisons.
